# Characterization of highly ferulate-tolerant *Acinetobacter baylyi* ADP1 isolates by a rapid reverse-engineering method

**DOI:** 10.1101/2021.09.07.459243

**Authors:** Jin Luo, Emily A. McIntyre, Stacy R. Bedore, Ville Santala, Ellen L. Neidle, Suvi Santala

## Abstract

Adaptive laboratory evolution (ALE) is a powerful approach for improving phenotypes of microbial hosts. Evolved strains typically contain numerous mutations that can be revealed by whole-genome sequencing. However, determining the contribution of specific mutations to new phenotypes is typically challenging and laborious. This task is complicated by factors such as the mutation type, the genomic context, and the interplay between different mutations. Here, a novel approach was developed to identify the significance of mutations in strains derived from *Acinetobacter baylyi* ADP1. This method, termed Rapid Advantageous Mutation ScrEening and Selection (RAMSES), was used to analyze mutants that emerged from stepwise adaptation to, and consumption of, high levels of ferulate, a common lignin-derived aromatic compound. After whole-genome sequence analysis, RAMSES allowed both rapid determination of effective mutations and seamless introduction of the beneficial mutations into the chromosomes of new strains with different genetic backgrounds. This simple approach to reverse-engineering exploits the natural competence and high recombination efficiency of ADP1. The growth advantage of transformants under selective pressure revealed key mutations in genes related to aromatic transport, including *hcaE*, *hcaK*, and *vanK*, and a gene, *ACIAD0482*, which is associated with lipopolysaccharide synthesis. This study provides insights into enhanced utilization of industrially relevant aromatic substrates and demonstrates the use of *A. baylyi* ADP1 as a convenient platform for strain development and evolution studies.

**Importance:** Microbial conversion of lignin-enriched streams is a promising approach for lignin valorization. However, the lignin-derived aromatic compounds are toxic to cells at relevant concentrations. Adaptive laboratory evolution is a powerful approach to develop more tolerant strains, but revealing the underlying mechanisms behind phenotypic improvement typically involves laborious processes. We employed *Acinetobacter baylyi* ADP1, an aromatic compound degrading strain that may be useful for biotechnology. The natural competence and high recombination efficiency of strain ADP1 can be exploited for critical applications such as the breakdown of lignin and plastics, abundant polymers composed of aromatic subunits. The natural transformability of this bacterium enabled us to develop a novel approach that allows rapid screening of advantageous mutations from ALE-derived aromatic-tolerant ADP1 strains. We clarified the mechanisms and genetic targets for improved tolerance towards common lignin-derived aromatic compounds. This study facilitates metabolic engineering for lignin valorization.

## 1. Introduction

The importance of adaptive laboratory evolution (ALE) (1, 2) in generating strains with desired traits is evidenced by success in improving the tolerance of production hosts towards stresses caused by non-optimal pH levels (3), high substrate or product concentrations (2, 4, 5), or other growth inhibitors (6, 7). Discovery of the associated genetic change can be accomplished by whole-genome sequencing (1, 2). However, it is challenging to determine the contribution of mutations, alone or in combination, to the evolved phenotype, as ALE typically yields multiple mutations (2). In addition, mutations may occur in poorly characterized genes. Some mutations may be neutral, others important for the evolutionary trajectory but not the final phenotype, and yet others may be deleterious hitchhikers (2).

Statistical methods have the potential to predict relevant mutations across a large number of independent ALE experiments (8), but a more profound understanding of the functional relevance of genetic change requires the reconstruction of strains with specific mutations (2). Such reconstruction, also referred to as reverse engineering, can be done by the introduction of selected mutations into reference strains, followed by comparative analyses (5, 7, 9–11). However, significant efforts may be required to synthesize alleles and integrate genetic changes in the appropriate location, especially when multiple mutations are tested. The bacterium used in our experiments, *Acinetobacter baylyi* ADP1 (hereafter ADP1), has unique advantages for determining the combinatorial effects of mutations. Its high efficiency of natural transformation and allelic replacement have long been exploited because linear DNA can be added directly to growing cultures. DNA responsible for a new phenotype is readily identified when it confers a growth advantage (12). As described here, we developed a high throughput method for the simultaneous analysis of multiple ALE-generated mutations.

ADP1 is an ideal model organism for biotechnology and synthetic biology (13) that has been used to study bacterial metabolism, engineering, and evolution (14–17). The potential of ADP1 as a production host for the synthesis of both native (18–22) and non-native products (20, 23, 24) has also been demonstrated. In our previous study, we engineered ADP1 for the production of 1- alkenes from ferulate (23), which represents one of the major compounds of alkaline-pretreated lignin (25–27). Engineering aromatic compound catabolism to valorize the lignin fraction of lignocellulose has important industrial potential (28, 29), and ADP1 is among the best bacterial candidates for this purpose (26). However, due to the inhibitory effects of lignin-derived aromatic compounds and the complexity of the associated pathways, natural pathways are not yet optimal for lignin valorization.

In this study, ALE was used to increase the tolerance and utilization of aromatic compounds to improve the suitability of ADP1 for these biotechnology applications. Catabolic pathways for aromatic compound degradation via the β-ketoadipate pathway in ADP1 and other bacteria have been well characterized (30–32). ALE has been shown to be effective in overcoming problematic aspects of complex regulation, transport, and enzyme specificity for aromatic compound degradation (17, 33). As described in this report, we characterized the phenotypes of different ALE lineages followed by whole-genome sequencing. Our new method, Rapid Advantageous Mutation ScrEening and Selection (RAMSES), facilitated the identification of the relevant mutations and the reverse engineering process. Finally, strains with increased tolerance were reconstructed by the new method and characterized. This study provides insights into enhancing the tolerance of production hosts towards lignin-derived aromatics and highlights the utility of ADP1.

## 2. Results

### 2.1. Adaptive laboratory evolution of *A. baylyi* ADP1 for high ferulate tolerance

Two parallel evolutions were previously carried out for ADP1 to improve the growth on ferulate as a sole carbon source (23), designated as G1 and G2 evolution lines here (Figure S1). Here, we carried out the ALE experiment also for two additional lineages to improve the tolerance towards ferulate, where 0.2% (w/v) casamino acids and 10 mM acetate were supplied along with ferulate, designated as T1 and T2 evolution lines (Figure S1). Acetate was added as an additional carbon source, as acetate is present in substantial amounts in the commonly used lignin-enriched stream (25, 26). This approach allows finding mutations that are potentially advantageous for both tolerance and utilization of aromatic compounds. To develop highly ferulate-tolerant strains, ferulate concentration was increased step-wise during the evolution experiments. A starting concentration of 55 mM was applied to the T1 and T2 evolution lines. The cells were daily passaged to fresh media for two months (360-375 generations), with the endpoint ferulate concentration being 135 mM for the T1/T2 evolution line. Individual isolates were obtained from end-point populations. Four isolates, which were named using ASA strain designations, were used for the subsequent studies (Figure S1): ASA500 and ASA501 were from G1 and G2 evolution lines respectively, and ASA502 and ASA503, were both from the T2 evolution line.

### 2.2. Characterization of the evolved strains

The evolved strains ASA500, ASA501, ASA502, and ASA503 were cultivated at different ferulate concentrations in 96-well plates, and wild-type ADP1, designated as WT ADP1, was used as the control. It was noted that a biphasic growth pattern was observed in some cases when cells were grown on the aromatic substrates (Figure S2). Therefore, to evaluate the tolerance of strains consistently and comprehensively at different conditions, we decided to use the time needed for cells to reach OD 0.8 as an indicator (later t_OD 0.8_). This indicator is influenced by both lag phase and growth rate.

All the evolved strains exhibited improved tolerance to ferulate, ASA500 had the best t_OD 0.8_ value (ASA500 and ASA501 Figure 1B, ASA502 and ASA503 Figure S3). In the presence of 20 mM ferulate, the t_OD 0.8_ was 42.5 h for WT ADP1, 12 h for ASA500, and 16 h for ASA501 (Figure 1B). For WT ADP1, a long lag phase accounted for most of t_OD 0.8_. When ferulate concentration was increased to 80 mM, the growth of WT ADP1 was completely inhibited while the t_OD 0.8_ value was prolonged to 17 h for ASA500, and 33.5 h for ASA501 (Figure 1B). A similar experimental set- up was employed to test the growth of ASA500 and ASA501 on *p*-coumarate and vanillate. *p*- coumarate, ferulate, and vanillate are all catabolized through the protocatechuate branch of the β- ketoadipate pathway. Vanillate is also an intermediate metabolite in the catabolism of ferulate (34) (Figure 1A). Improved tolerance towards *p*-coumarate and vanillate was also observed from the two isolates ASA500 and ASA501 (Figure 1B). Vanillate seemed to be less toxic than ferulate and *p*-coumarate, as indicated by the growth of WT ADP1 on this substrate.

**Figure 1.**
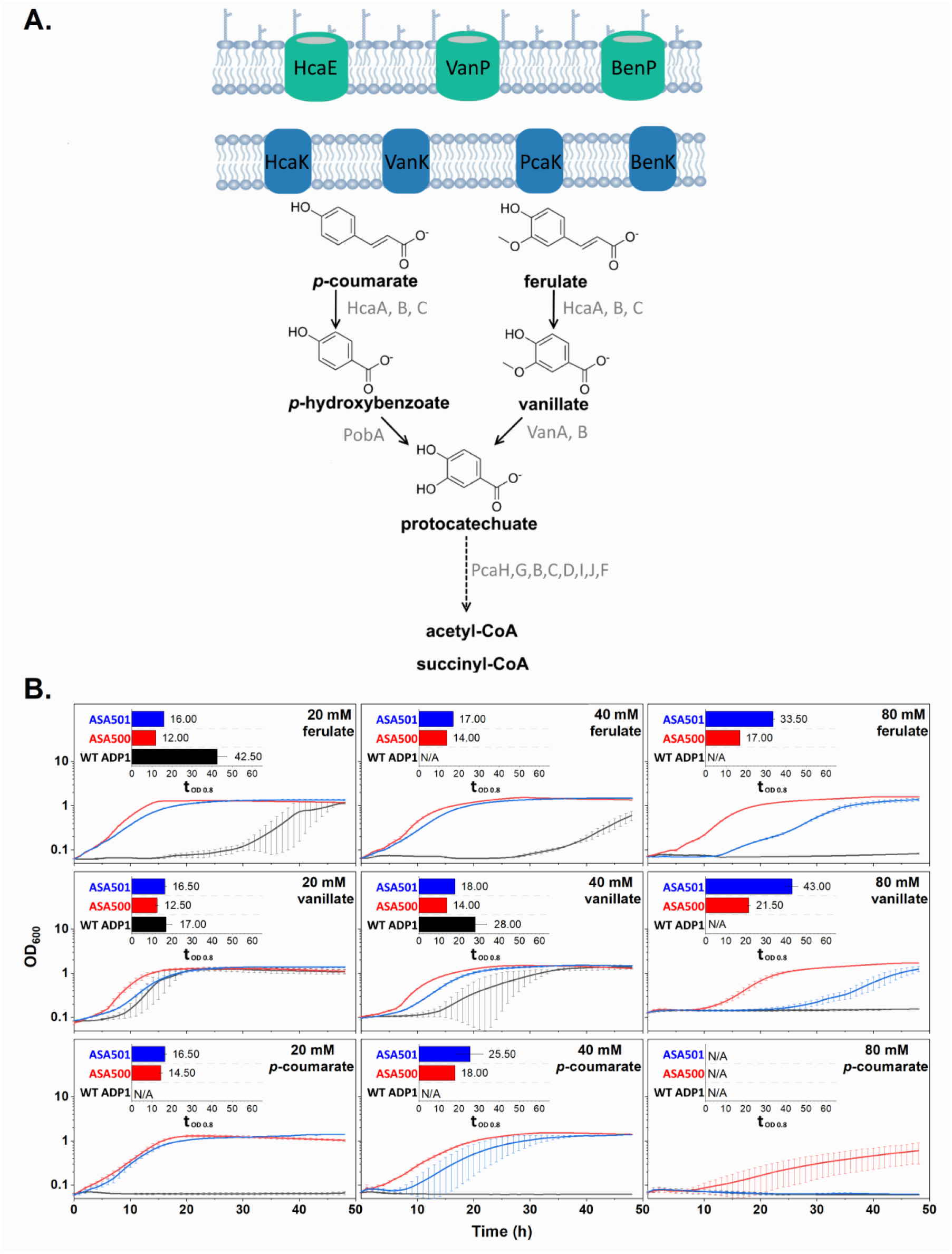
Growth of ASA500, ASA501, and WT ADP1 on ferulate, vanillate, and *p*-coumarate. (A) Possible transport system (porins colored in green and transporters colored in blue) for aromatic acids and the β-ketoadipate pathway in ADP1. The pathway indicated by the dashed arrow involves multiple steps. (B) Growth of the strains ASA500, ASA501, and WT ADP1 at different concentrations of ferulate, vanillate, and *p*-coumarate. All the strains were cultivated in mineral salts media with the corresponding aromatic compound as the sole carbon source. The time needed for the cells to reach the OD of 0.8 was used as the indicator to evaluate the tolerance (t_OD 0.8_). The indicator is calculated only when both replicates reached OD 0.8 within 48 h. All the values were calculated from two replicates and the error bars indicate the standard deviations. The y-axis is shown in log10 scale.

Although ASA502 and ASA503 evolved in the presence of 0.2% casamino acids and 10 mM acetate, both showed improved growth in ferulate as the sole carbon source (Figure S3). The two strains were further cultivated in elevated ferulate concentrations while being supplemented with casamino acids and acetate. When the ferulate concentration was increased from 40 mM to 80 mM, there was a 5 h of increase in the t_OD 0.8_ for both strains (Figure S4), which was shorter than the greater than 10 h increase observed when ferulate was the sole carbon source (Figure S3). In 40 mM ferulate, WT ADP1 showed diauxic growth characteristic of the sequential consumption of carbon sources; the aromatic degradative pathway is known to be repressed in the presence of acetate through catabolite repression (31). Diauxic growth was not observed for ASA502 and ASA503. However, HPLC analysis showed that while acetate and ferulate were consumed sequentially when both substrates were present, the ferulate was rapidly consumed after acetate was depleted (data not shown). Interestingly, an increase of acetate concentration from 10 mM to 50 mM improved the tolerance of WT ADP1 towards ferulate (Figure S4).

The tolerance towards aromatic acids may also be affected by the pH of media, possibly related to the protonation of the acids. Protonated aromatic acids can passively diffuse across bacterial cell membranes (35). The results in a Supplemental Note demonstrated an improved growth of WT ADP1 on ferulate in higher pH, which favors deprotonation of weak acids.

### 2.3. Genome sequencing of the evolved strains

Whole genome sequencing was performed to discover the mutations in the evolved isolates. The sequencing reads from the five sequenced strains (ASA500, ASA501, ASA502, ASA503, and WT ADP1) were mapped to the reference genome of *A. baylyi* ADP1 (GenBank: CR543861). As some sequence variants that differ from the reference genome are present in the parent strain WT ADP1 (Table S3), these variants were subtracted from the mutation pool of the evolved isolates. All the mutations in the evolved isolates are summarized in Table 1. The number of mutations in coding and non-coding regions for each strain is summarized in Figure 2. As ASA502 and ASA503 were isolated from the same population, they shared several mutations. For all the evolved strains, many of the mutations were found in the genes whose products are membrane proteins or involved in cell envelope modification. Notably, some of these genes are associated with aromatic transport, including *hcaE*, *vanK*, and *hcaK* (34, 36, 37). The gene *hcaE* encodes an outer membrane porin, and both *vanK* and *hcaK* encode transport proteins belonging to the major facilitator superfamily. The gene *hcaE* was mutated in all the four strains: insertion of the IS (insertion sequence) element, IS*1236*, for ASA500 and ASA501, and a single nucleotide insertion for ASA502 and ASA503.

**Figure 2.**
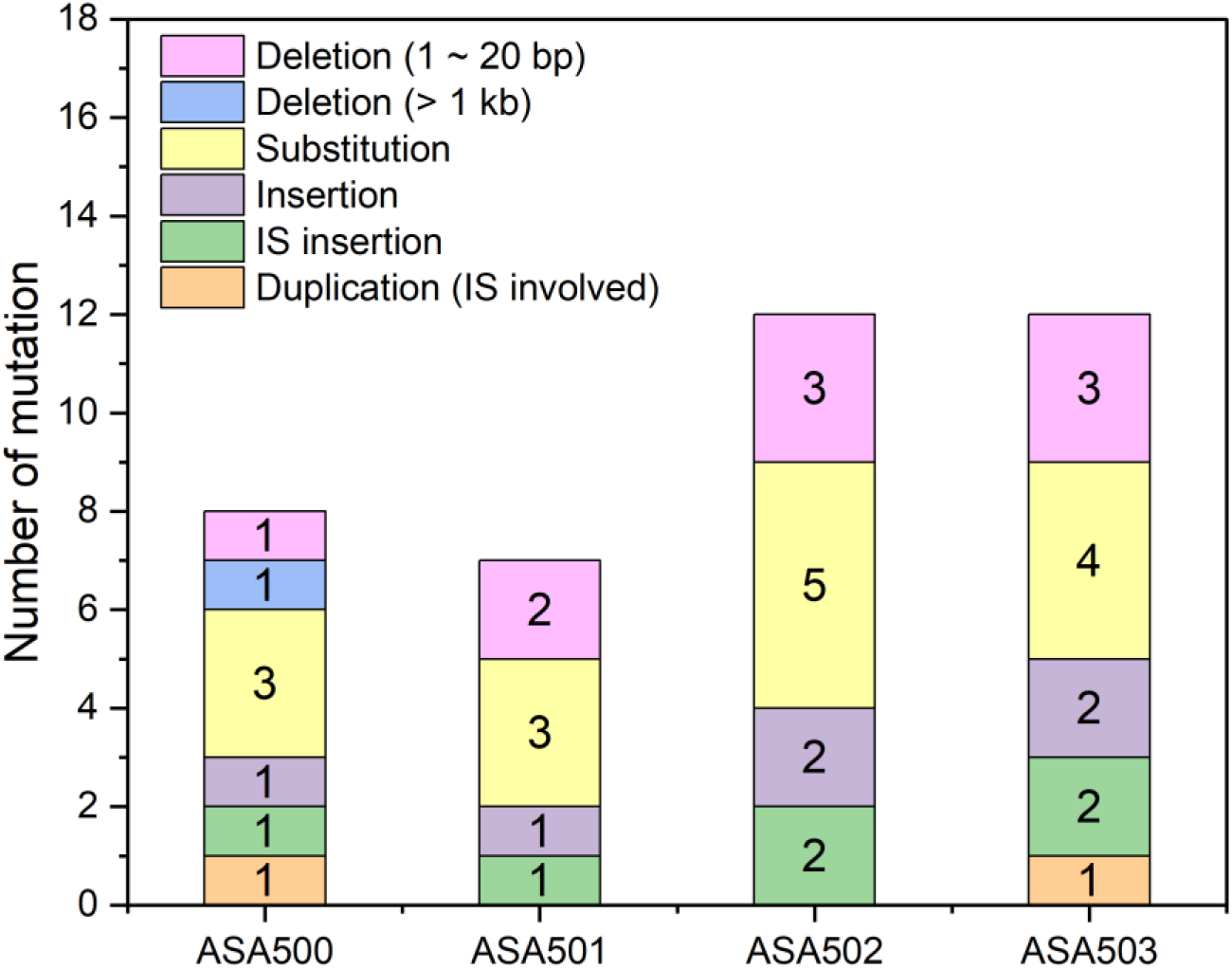
Number of mutations in the genomes of each evolved strain.

**Table 1.** Mutations in the evolved isolates.

| Gene locus ID (name) <sup>a</sup> | Position <sup>a</sup> | Description <sup>a</sup> | Mutation type | DNA change | Protein effect | ASA 500 | ASA 501 | ASA 502 | ASA 503 |
| --- | --- | --- | --- | --- | --- | --- | --- | --- | --- |
| *ACIAD1702* (*pcaU*) | 1708197 | Regulatory protein for *pca* operon (activator) | Substitution (transition) | G → A | P250L (CCA→CTA) | + | + |  |  |
| *ACIAD1722* (*hcaE*) | 1730279-1730280 | Porin | Insertion (tandem repeat) | (G)3 → (G)4 | Frameshift |  |  | + | + |
| *ACIAD1722* (*hcaE*) | 1730384-1730385 | Porin | Insertion (IS element) | +IS, +AGG | Frameshift | + | + |  |  |
| *ACIAD1727* (*hcaK*) | 1736544-1736548 or 1736548-1736552 | Transporter | Deletion | -TGCTG or -GTGCT | Frameshift* |  | + |  |  |
| *ACIAD0982* (*vanK*) | 967651-968787 | Transporter | Deletion (> 1kb) | 1137 bp deletion | Loss of function | + |  |  |  |
| *ACIAD2867* | 2807553 | Putative Na<sup>+</sup>/H<sup>+</sup> antiporter | Substitution (transition) | G → A | A247V (GCA→GTA) | + |  |  |  |
| *ACIAD2265* | 2236575-2236576 | Putative rare lipoprotein A family | Insertion | +C | Frameshift |  | + |  |  |
| *ACIAD2365* (*msbA*) | 2322575 | Lipid transport protein | Substitution (transition) | G → A | A486V (GCG→GTG) |  |  | + | + |
| *ACIAD0648* (*secA*) | 639202 | Preprotein translocase | Substitution (transition) | C → T | P796L (CCA→CTA) |  |  | + |  |
| *ACIAD0482* | 476258-476269 or 476264-476275 | putative glycosyltransferase | Deletion | -TGAGGAAAGGCT or -AAGGCTTGAGGA | Δ166-169 | + |  |  |  |
| *ACIAD0602* | 591935 | Putative glycosyltransferase | Deletion (tandem repeat) | (T)5 → (T)4 | Frameshift |  |  | + | + |
| *ACIAD1405* (*iscR*) | 1399615 | Repressor of the *iscRSUA* operon | Substitution (transition) | G → A | Truncation |  |  | + | + |
| *ACIAD0260* (*gacA*) | 261189 | Response regulator | Deletion | -A | Frameshift |  |  | + | + |
| *ACIAD3465* | 3392835 | Putative two-component sensor | Substitution (transversion) | G → T | G881C (GGT→TGT) |  | + |  |  |
| *ACIAD2274* (*sthA*) | 2243241 | Soluble pyridine nucleotide transhydrogenase | Substitution (transition) | G → A | A462V (GCT→GTT) |  |  | + | + |
| *ACIAD0438* (*rne*) | 436298 | Ribonuclease E | Deletion | -A | Frameshift |  | + |  |  |
| *ACIAD3194* (*rpoA*) | 3122059 | RNA polymerase alpha subunit | Substitution (transversion) | G → T | P291Q (CCA→CAA) |  | + |  |  |
| *ACIAD0308* (*rpoC*) | 307439 | RNA polymerase beta subunit | Substitution (transition) | A → G | D285G (GAT→GGT) |  |  | + |  |
| *ACIAD1220* | 1223930 | Conserved hypothetical protein | Substitution (transversion) | C → G | G84A (GGA→GCA) | + |  |  |  |
| *ACIAD3457* to *ACIAD3481* | 3380313-3408297 | N/A | Duplication (IS involved) | Duplication | N/A | + |  |  |  |
| *ACIAD3459* to *ACIAD3486* | 3380938-3413938 | N/A | Duplication (IS involved) | Duplication | N/A |  |  |  | + |
| N/A | 2808313-2808314 | Non-coding region 21 bp upstream of *ACIAD2867* | Insertion (IS element) | +IS, +AAC | N/A |  |  | + | + |
| N/A | 474157-474158 | Non-coding region 135 bp upstream of *ACIAD0481* | Insertion (IS element) | +IS, +TGT | N/A |  |  | + | + |
| N/A | 2766926-2766927 | Non-coding region between *ACIAD2829* and *ACIAD2832* | Insertion (tandem repeat) | (T)6 → (T)7 | N/A | + |  | + | + |
| N/A | 2236147 | Non-coding region between *ACIAD2264* (*mltB*) and *ACIAD2265* | Deletion (tandem repeat) | (T)7 → (T)6 | N/A |  |  | + | + |
| N/A | 967592 | Non-coding region 17 bp upstream of *ACIAD0980* (*vanA*) | Substitution (transition) | C → T | N/A |  |  |  | + |

\* For HcaK, the UniProt entry Q7X0E0 (with 410 residues) was used as a reference to evaluate the protein effect.
<sup>a</sup> Locus IDs, mutation positions, and descriptions were assigned according to the reference genome CR543861 (GeneBank entry).

An 1137 bp deletion extending from the position 732 bp upstream of *vanK* to its CDS position 405 was identified in ASA500, and a 5 bp deletion in *hcaK* was identified in ASA501. All the mutations in the three genes would likely result in loss of protein function.

We analyzed the emergence of the IS insertion in *hcaE* and the 1137 bp deletion in *vanK* by PCR-amplification of the target regions from the genomes of samples from the evolving populations taken at different times during the experiments. It was found that the *hcaE* mutation had already emerged on day 3 (≈11 generations, ferulate concentration = 45 mM) for both G1 and G2 evolution lines (Figure 3A). Considering that the *hcaE* mutations in the two independently evolved strains, ASA500 and ASA501, are identical, the *hcaE* mutations were probably from the same origin and had already occurred in the pre-culture stage where 45 mM ferulate was applied (Figure S1). The deletion in the *vanK* region had already occurred in the G1 evolution line on day 20 (≈119 generations, ferulate concentration = 80 mM) and had been fixed between day 40 (≈236 generations, ferulate concentration = 115 mM) and day 50 (≈299 generations, ferulate concentration = 120 mM) (Figure 3B).

**Figure 3.**
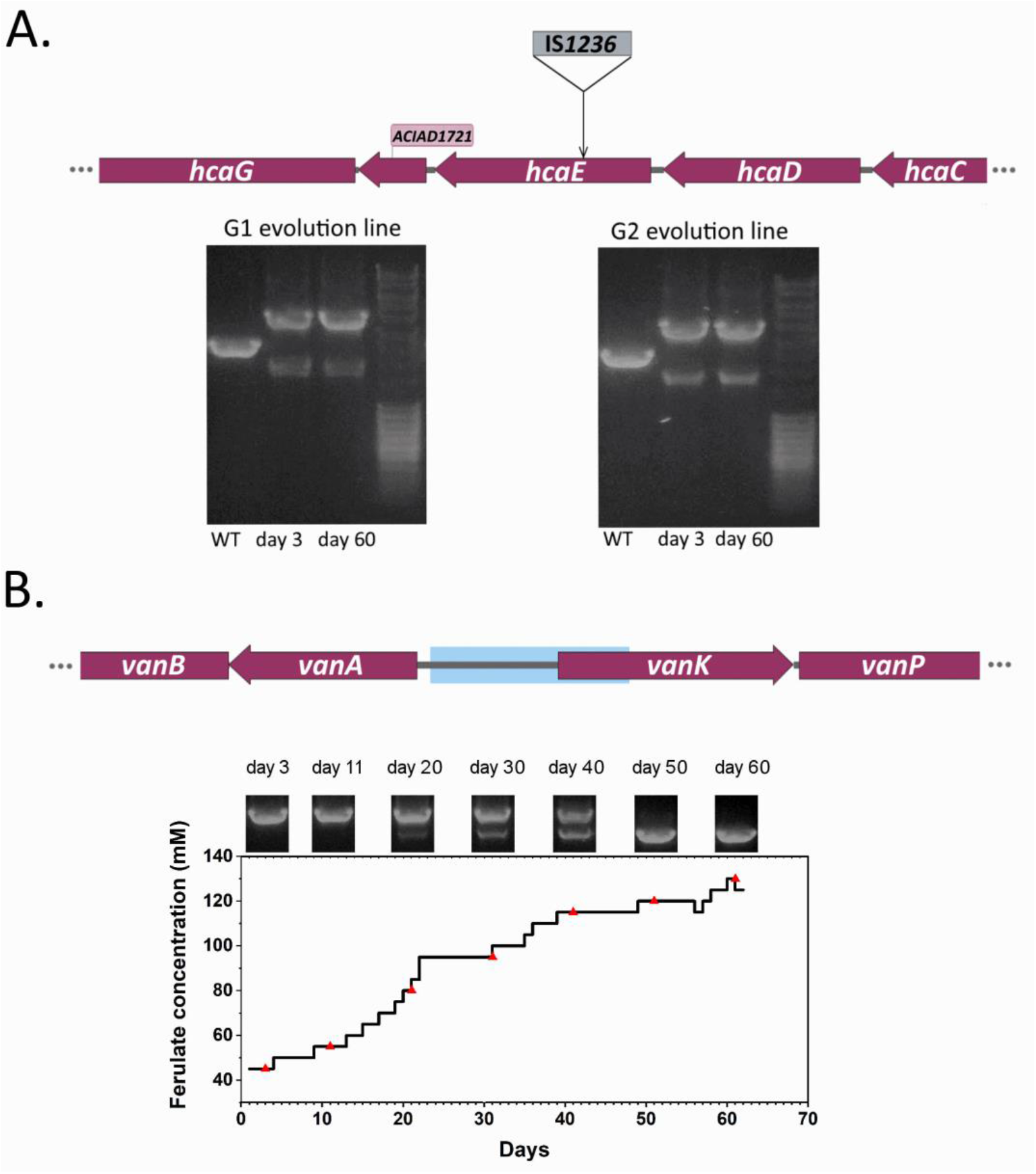
Temporal occurrence of the mutations in *hcaE* and *vanK* in G1 and G2 evolution populations. (A) The *hcaE* mutation caused by IS*1236* insertion and gel electrophoresis showing the genotypic change in *hcaE* in G1 and G2 evolution populations sampled on different days. (B) Deletion in the *vanK* region and the genotypic change of this region in G1 evolution populations over time. The deletion of 1137 bp (highlighted in blue) extends from the upstream of *vanK* to its CDS position of nucleotide 405.

Other mutations that were likely to cause loss of function were found in the genes *ACIAD2265*, *iscR*, *gacA*, and *ACIAD0602* (Table 1). *ACIAD2265,* which was mutated in ASA501, is predicted to encode a lytic transglycosylase involved in cell wall organization. The other genes, *iscR*, *gacA*, and *ACIAD0602*, were found to be mutated in ASA502 and ASA503. *iscR* potentially encodes a repressor of the *iscRSUA* operon, which is involved in the assembly of Fe-S clusters. Fe-S clusters are important in enzymes for aromatic compound degradation; for example, they act as co-factors of a two-component vanillate demethylase (VanAB) for the conversion of vanillate into protocatechuate (38). The gene *gacA* encodes a response regulator whose deletion has been characterized in *A. baumannii* and would lead to up/down-regulation of a large number of genes (39). Interestingly, further analysis of the up/down-regulated genes showed that some genes are related to aromatic catabolism and uptake. *ACIAD0602* encodes a putative glycosyltransferase which shares >80% identity with GtrOC4 in *A. baumannii* by NCBI protein blast. GtrOC4 was proposed to be involved in the outer core synthesis of lipo-oligosaccharides (40).

Besides the aforementioned IS*1236* insertion in *hcaE* in ASA500 and ASA501, IS*1236* insertion was also identified in two non-coding regions in ASA502 and ASA503 (Table 1): one is 21 bp upstream of *ACIAD2867*, and another one is 135 bp upstream of *ACIAD0481*. Consistent with the previous report (41), all the IS*1236* insertions generated a small duplication, which resulted in 3 bp repeats flanking the inserted IS element, as is known to occur for the mechanism of its transposition.

### 2.4. IS-involved gene duplication was identified in the evolved strains

Gene duplication was found in the evolved strains ASA500 and ASA503, which was probably mediated by IS*1236*. Sequencing analysis showed that the strains ASA500 and ASA503 had DNA regions at similar genomic positions with sequencing coverages 2-fold higher than those of the genomes (Figure S5 and Table S4), suggesting a duplication event. The region in ASA500 had a size of approximately 28 kb covering the whole coding sequences of the genes from *ACIAD3457* to *ACIAD3481*, while the region in ASA503 had a size of about 33 kb extending from *ACIAD3459* to *ACIAD3486* (Table S4). Most of the involved genes were shared by the duplicated regions in ASA500 and ASA503. The sequence of IS*1236* was found at each junction of the region (Figure S6), suggesting that the duplicated region was flanked by IS*1236*. A potential explanation is that the region could be duplicated in the form of a composite transposon that might be inserted in one of the original IS*1236* sites. The duplications in the two strains were not identical, indicating that they resulted from independent events. However, the duplication was absent in ASA502, which was isolated from the same population as ASA503.

### 2.5 Rapid selection of advantageous mutations and reverse engineering

To select and reverse-engineer the mutations that confer significantly improved tolerance towards ferulate, a novel approach (Rapid Advantageous Mutation ScrEening and Selection, RAMSES, Figure 4) was implemented. This method is based on ADP1’s natural transformation, active homologous recombination machinery, and efficient enrichment of advantageous mutants under selective conditions. As illustrated in figure 4, the transformation is done by simply adding the amplified mutated alleles containing flanking regions of sequence identity to the chromosome to the cell culture (for liquid medium transformation) or the colony (for solid medium transformation). The cultures after transformation are then used to inoculate different selective media with incremental selective pressures (here aromatic concentration). The use of a range of selective pressures enables finding a suitable condition for selection. The advantageous mutations are selected if the cells transformed with the corresponding alleles show significantly improved growth under the conditions used for growth.

**Figure 4.**
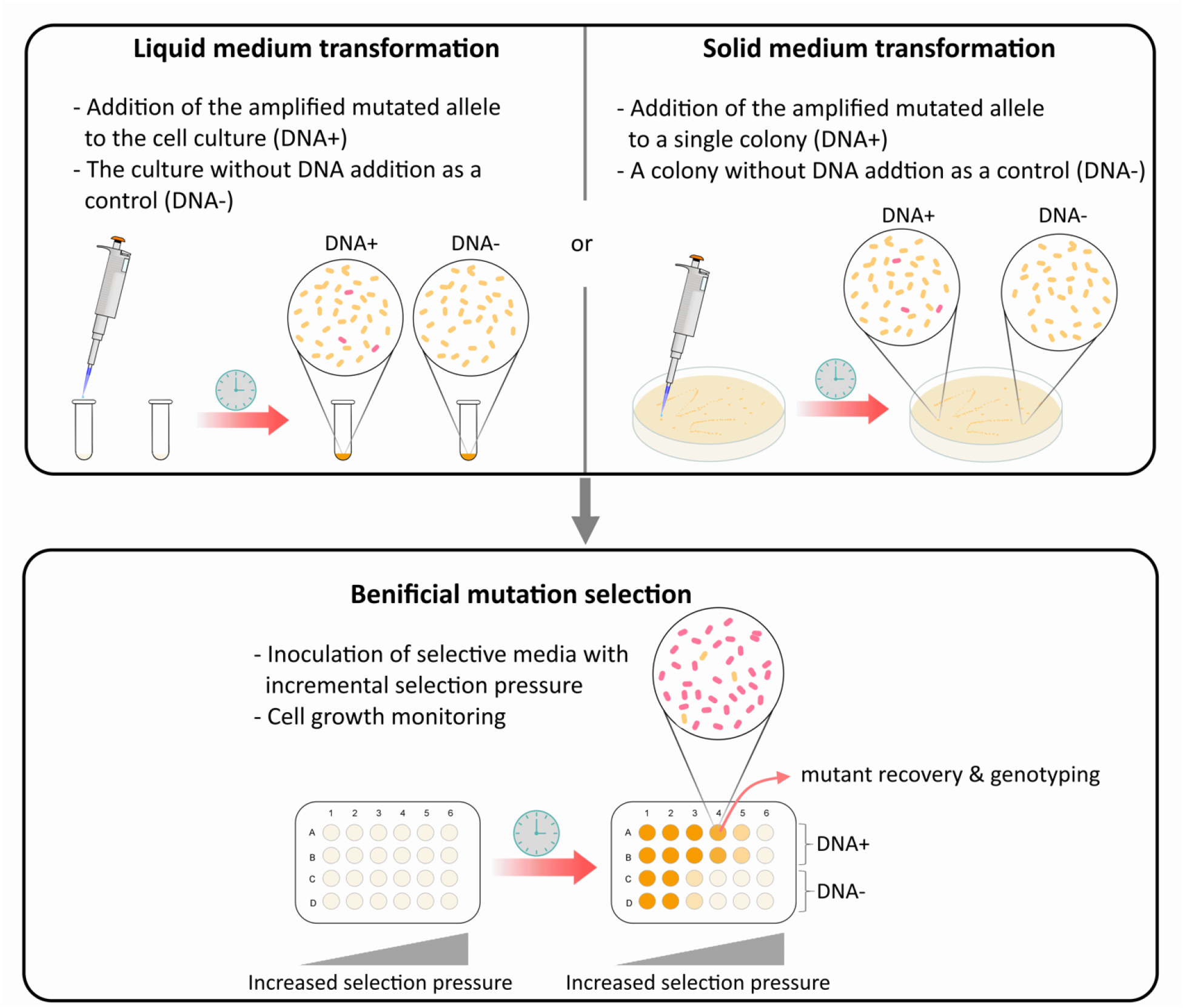
Schematic of RAMSES. The mutated allele amplified from the evolved strain is used to transform the background strain by direct addition of the purified PCR product to the exponentially growing cells (in small volume) or newly emerging colony. The linear DNA is incorporated into the chromosome by allelic replacement. The cultures after transformation are used to inoculate different media with incremental selective pressures (here aromatic concentration) in a multi-well plate. The growth is monitored based on the measurement of optical density. The mutants (colored in red) containing the advantageous mutation can grow robustly and get enriched at the level of selective pressure such that it outcompetes the background strain, resulting in a different growth profile compared to the control. The mutant can be recovered from the culture by an additional enrichment step under the same (or higher) selective pressure and subsequent isolation, and the corresponding mutation(s) can be confirmed by sequencing (or PCR if possible).

RAMSES was performed with seven mutated alleles from the evolved isolates ASA500 and ASA501, including the *pcaU500* (P250L), *hcaE500* (frameshift by IS insertion), *vanK500* (loss of function by deletion), *hcaK501* (frameshift), *ACIAD2867_500* (A247V), *ACIAD2265_501* (frameshift), and *ACIAD0482_500* (Δ166-169 in amino acid sequence). The transposon-free *A. baylyi* ADP1 (42), designated as ISx, was used as the background strain for the RAMSES. The strain, in which all the six copies of IS*1236* were deleted, has been shown to exhibit a more stable phenotype and increased transformability (42). The transformation was performed in liquid medium. The transformed cells were transferred to the selective media containing 20 mM, 40 mM, 60 mM, and 80 mM ferulate. At 60 mM of ferulate, the cells transformed with *hcaE500* and *hcaK501* showed evident benefit, and their growth curves could be clearly distinguished from those of the controls (Figure 5), indicating the significance of the *hcaE* and *hcaK* mutations. The benefit from the mutation in *hcaE* is consistent with the observation that *hcaE* was mutated in all four evolved isolates. The other mutated alleles, such as the *pcaU500* and *ACIAD0482_500* mutations, showed detectable but less prominent effects or large variances between replicates.

**Figure 5.**
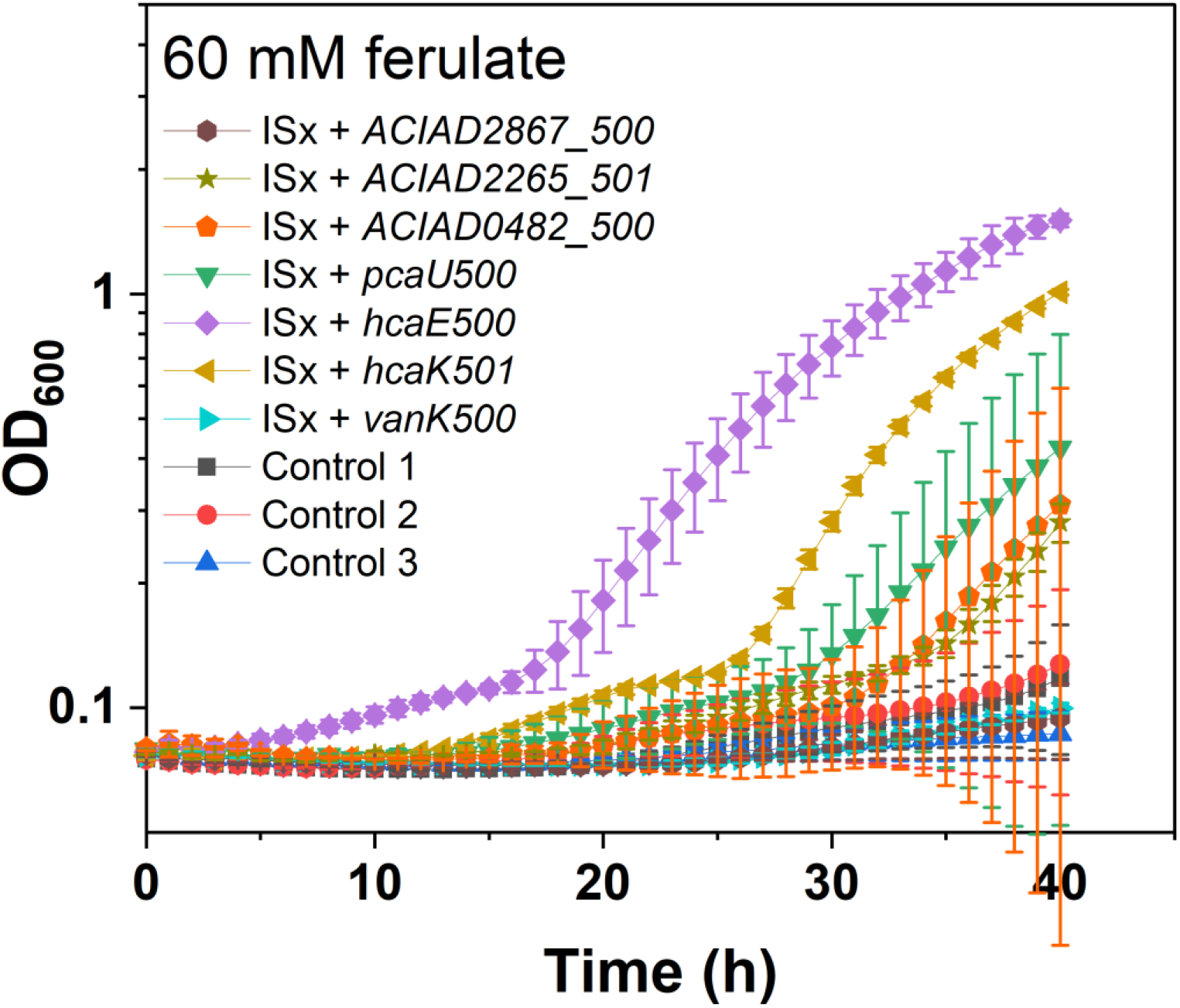
Initial screening of advantageous mutations by RAMSES. Growth of ISx in 60 mM ferulate after being transformed with the selected mutated alleles is shown. Control 1: ISx without any treatment. Control 2: ISx treated with water. Control 3: ISx treated with a non-mutated allele. The OD values were calculated from two replicates. The error bars indicate the standard deviations. The y-axis is shown in log10 scale.

Although the mutation-transformed cells could be directly isolated from the initial screening experiment, we confirmed the reproducibility of the method by re-introducing the mutations to the ISx strain by solid medium transformation. As the *hcaE* mutation found in ASA500/ASA501 occurred in the early stage of the G evolution lines (Figure 3A), it was chosen as the first mutation to be introduced into ISx. ISx showed improved growth at the elevated ferulate concentration (20 mM) after being transformed with *hcaE500* (Figure 6A). An additional round of cultivation under the selection condition was performed to further enrich the *hcaE* mutant. PCR analysis from the genome of the enriched population showed the existence of both wild-type and the mutated *hcaE* genotypes (Figure 6A), indicating the enrichment of the *hcaE* mutant. The pure strain containing the mutant *hcaE* was further isolated and designated as ASA504. The mutated allele *vanK500* was also chosen to transform ISx, given its propagation in the G1 evolution population over time (Figure 3B) and the role of the gene related to aromatic transport (36). However, only one of the two replicate populations that were transformed with *vanK500* showed improved growth and enrichment of the *vanK* mutant (Figure S7). The pure strain containing *vanK500* was further isolated and designated as ASA505.

**Figure 6.**
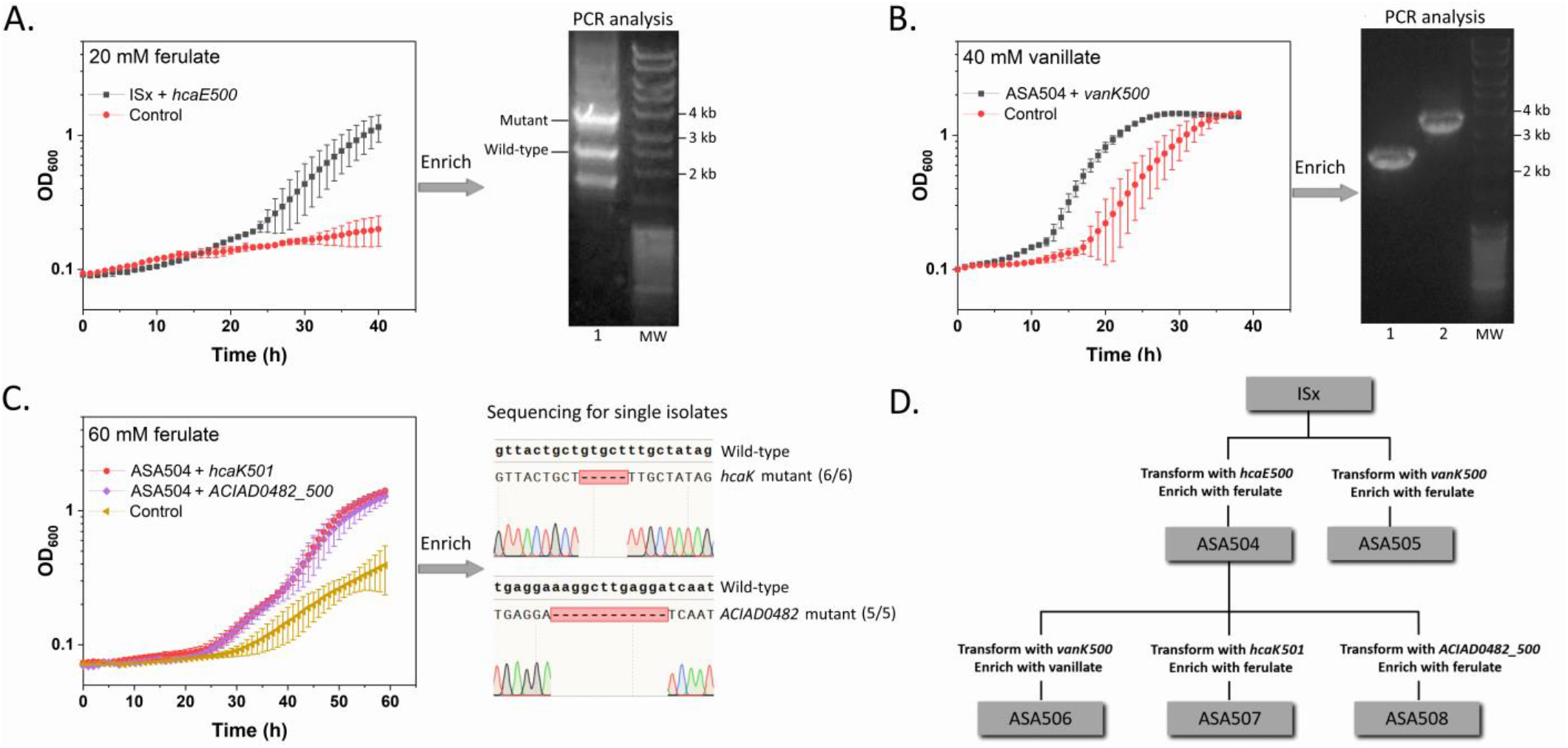
Reverse engineering of the selected mutations by RAMSES. (A) Reverse engineering of *hcaE500* into ISx. Growth of ISx in 20 mM ferulate after transformation is shown. Control: ISx without transformation. To analyze *hcaE*, PCR analysis was performed from the genome of the transformed population after further enrichment. Lane 1: the transformed population. (B) Reverse engineering of *vanK*500 into ASA504 (reconstructed mutant *hcaE*). Growth of ASA504 in 40 mM vanillate after transformation is shown. Control: ASA504 without transformation. To analyze *vanK*, PCR analysis was performed from the genome of the transformed population after further enrichment. Lane 1: the transformed population. Lane 2: ASA504 containing wild-type *vanK*. (C) Reverse engineering of *hcaK501* and *ACIAD0482_500* into ASA504 (reconstructed mutant *hcaE*). Growth of ASA504 in 60 mM ferulate after transformation is shown. Control: ASA504 without transformation. The genes *hcaK* and *ACIAD0482* of the single isolates from the enriched populations were analyzed by sequencing. Pure mutant strains were recovered by further isolation. (D) The reconstructed strains derived from ISx. The OD values were calculated from two replicates. The error bars indicate the standard deviations. The y-axis is shown in log10 scale.

We next used ASA504 (reconstructed mutant *hcaE*) as the parent strain for the introduction of other mutated alleles, including *vanK500*, *hcaK501*, *pcaU500*, and *ACIAD0482_500*. The ASA504 populations transformed with *vanK500* and *pcaU500* respectively did not show significantly improved growth at the ferulate concentrations tested (20-80 mM) (data not shown). However, it was found that ASA504 had poor growth on vanillate, while ASA505 (reconstructed mutant *vanK*) had improved growth in the same condition (Figure S8). Therefore, we hypothesized that *vanK500* would restore the growth of ASA504 on vanillate. Next, we transformed ASA504 with *vanK500* and used vanillate as the selective pressure for mutant enrichment. As expected, the population of ASA504 showed improved growth on vanillate after being provided with the *vanK500* (Figure 6B). PCR analysis of *vanK* from the genomic DNA extracted from the enriched population showed only the band of *vanK500* (Figure 6B), indicating the predominance of this allele. A streak-purified isolate was designated as ASA506. Transforming ASA504 with *hcaK501* and *ACIAD0482_500* led to improved growth at 60 mM of ferulate (Figure 6C). After further enrichment with the same selective pressure, pure isolates were obtained from each of the populations. Six isolates from the *hcaK501* transformed-population and five isolates from the *ACIAD0482_500* transformed-population were analyzed by Sanger sequencing for *hcaK* and *ACIAD0482* respectively. All these isolates were shown to contain the corresponding mutated alleles (Figure 6C). The mutation in *hcaK501* would result in a frameshift (based on the HcaK sequence from UniProt entry Q7X0E0) and likely caused loss of protein function, while the mutation in *ACIAD0482_500* would lead to deletion of 4 amino acids. The resulting mutants are designated as ASA507 and ASA508 respectively. All the reconstructed strains were summarized in Figure 6D.

### 2.6. Characterization of the reconstructed strains

To compare the growth on ferulate between the reconstructed strains, they were cultivated at different ferulate concentrations. WT ADP1, ISx, ASA500 (evolved isolate), and ASA501 (evolved isolate) were also cultivated for comparison. Although WT ADP1 seems to differ from ISx only in the copy number of the IS*1236* element, it had a better growth than ISx at 20 mM ferulate (Figure 7). Compared to the reference strain ISx, the single mutants, ASA504 (reconstructed mutant *hcaE*) and ASA505 (reconstructed mutant *vanK*), showed improved growth at 20 mM of ferulate (Figure 7). Both strains exhibited improved tolerance also towards *p*- coumarate (Figure S8). However, the growth of the two single mutants was strongly inhibited at 40 mM ferulate. Introduction of *vanK500* only slightly improved the growth of ASA504, as indicated by the growth of ASA506 (reconstructed mutant *hcaE* and *vanK*) (Figure 7). This result was consistent with the previous failed attempt to enrich the *hcaE* and *vanK* double mutant using ferulate. However, the growth of ASA504 on vanillate was significantly improved by introducing *vanK500* (Figure S8). The genes *vanK* and *vanP* may be under the control of the same promoter due to their proximity. The mutation in *vanK500* would likely cause loss of *vanK* promoter region, which may negatively affect the expression of the downstream gene *vanP*. It was further explored (see Supplemental Note) whether deletion of both *vanK* and *vanP,* or *vanP* alone had the same effect as the mutation in *vanK500* in terms of improving the tolerance of ASA504 to vanillate. Both the *hcaK* and the *ACIAD0482* mutations further improved the tolerance of ASA504 to ferulate, as indicated by the robust growth of ASA507 (reconstructed mutant *hcaE* and *hcaK*) and ASA508 (reconstructed mutant *hcaE* and *ACIAD0482*) at 40 mM (Figure 7). ASA507 had the best growth on ferulate among the reconstructed strains, as it was the only reconstructed strain that grew robustly at 60 mM ferulate. Although a direct comparison between the evolved strain ASA500/ASA501 and ASA507 is complicated by their derivations from different parent strains, it is clear that there is potential to further recapitulate the evolved tolerance patterns by introducing other mutations.

**Figure 7.**
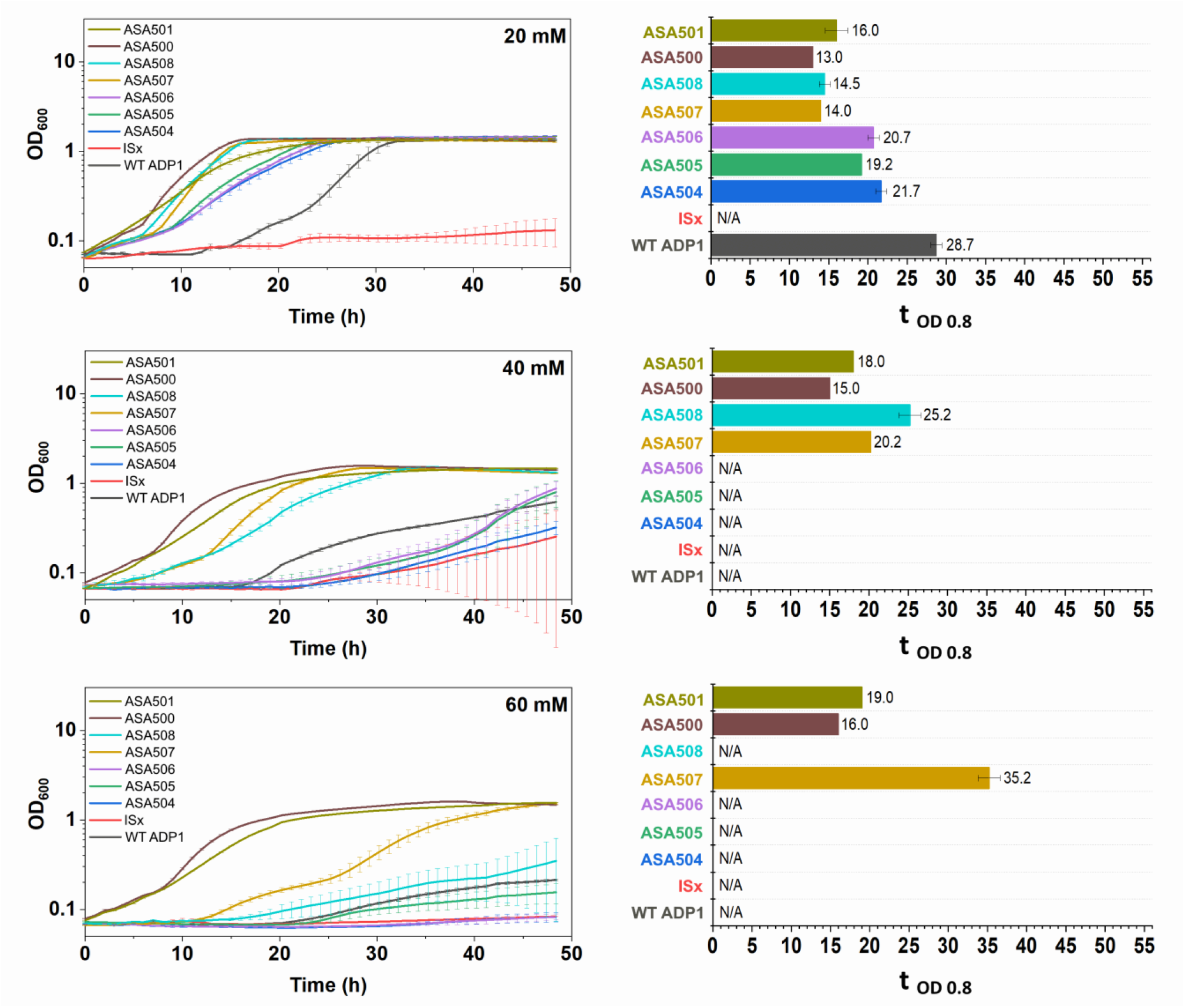
Growth comparison between the reconstructed strains at different ferulate concentrations (20 mM, 40 mM, and 60 mM). ASA500 and ASA 501 are the evolved strains, and ASA504-508 are the reconstructed strains. All the strains were cultivated in mineral salts media with the ferulate as the sole carbon source. The time needed for the cells to reach the OD of 0.8 was used as the indicator to evaluate the tolerance (t _OD 0.8_). The indicator was calculated only when both replicates reached OD 0.8 within 48 h. All the values were calculated from two replicates and the error bars indicate the standard deviations. The y-axis is shown in log10 scale

## 3. Discussion

Lignocellulose biorefining has been demonstrated as a sustainable method for the production of fuels, chemicals, and materials (43). To date, the polysaccharide fraction of lignocellulose is of primary interest for the downstream conversion, whereas the lignin fraction is usually regarded as a waste, a low value-added product, or a source for process heat. Recent analysis has indicated that lignin valorization is essential to increase the sustainability and economic viability of lignocellulose-based industries (44, 45). The success of lignin valorization largely relies on lignin depolymerization and subsequent upgrading of the heterogeneous lignin-derived aromatics (44). However, due to the heterogeneity and impurity of lignin, it is a very challenging feedstock for chemical processes (29). The microbial valorization of lignin has been suggested (25, 27, 29); the aromatic catabolic pathways in some microbes allow the “funneling” of various aromatic species into two key aromatic ring-cleavage substrates, commonly protocatechuate and catechol, which can be further channeled to central carbon metabolism. Salvachúa *et al.* examined fourteen bacteria for their ability to utilize a lignin-enriched substrate (26); *A*. *baylyi* ADP1 was the best among the tested bacteria both to degrade high molecular weight lignin and consume low molecular weight lignin-derived compounds. The use of *A*. *baylyi* ADP1 as a host of engineering for biological lignin valorization is further warranted by its straightforward genome editing (14, 46) and the rapidly increasing number of available genetic tools (46).

Apart from the availability of lignin-derived aromatics for use as a substrate, their toxicity must be considered, which is regarded as a major challenge in the biological upgrading of lignin-enriched streams (47, 48). All the aromatic compounds tested in the current study showed varying degrees of toxicity when used as sole carbon sources: the growth of wild-type ADP1 on ferulate was significantly impaired when the concentration was increased from 20 mM to 40 mM.

Moreover, 80 mM was lethal to the cells, and cell growth was not observed in 20 mM *p*-coumarate (Figure 1B). In a previous study, a 33% reduction of growth rate was observed in glucose-grown *Pseudomonas putida* KT2440 and *Escherichia coli* MG1655 in the presence of 61 mM and 30 mM *p*-coumarate respectively (49). Consequently, the use of batch fermentation is greatly limited, and suitable fed-batch strategies need to be developed for substrate feeding without reaching toxicity limits. Some aromatics can cause severe growth impairment at much lower concentrations. For example, benzoate and catechol have been reported to completely inhibit glucose-grown *P*. *putida* KT2440 at 50 mM and 8 mM respectively (47, 50). In addition, prolonged contact with toxic aromatic compounds, even at low concentrations, may lead to other cellular malfunctions, such as cellular energy shortage, as demonstrated by Kohlstedt *et al.* (47). Biotransformation can become more challenging when the lignin-derived aromatics serve as the sole carbon sources, especially if the products of interest require high levels of carbon substrate to sustain the synthesis of desired products.

In our attempts to discover the tolerance mechanisms behind the evolved strains, aromatic-specific transport was found to play an important role. The loss-of-function mutations in the genes *hcaE*, *hcaK*, and *vanK* were identified to be advantageous by RAMSES. Reconstruction of the *hcaE* mutation improved the tolerance towards both ferulate and *p*-coumarate. This gene encodes an outer membrane porin, and it clusters with other genes responsible for ferulate and *p*-coumarate catabolism (34), implying that the porin may act on hydroxycinnamates. Interestingly, the *hcaE* mutation resulted in decreased tolerance towards vanillate. HcaE might have a low specificity for vanillate. The tolerance towards vanillate was improved by reconstruction of the *vanK* mutation. The gene *vanK* encodes a transporter belonging to the major facilitator superfamily. Its location near the *vanAB* operon, which is responsible for vanillate catabolism, implies that vanillate may be transported by VanK, as proposed previously (51). VanK has been also reported to mediate the uptake of two other intermediates in the aromatic catabolic pathway, protocatechuate and *p*- hydroxybenzoate (36). The combination of the *hcaE* and *hcaK* mutations further improved the tolerance to ferulate. The gene *hcaK*, transcribed in the opposite direction of the *hca* operon by a bidirectional promoter (52), encodes a transporter which also belongs to the major facilitator superfamily. It is possible that ferulate and *p*-coumarate are both transported by HcaK. Loss of function mutations in these genes related to aromatic acid transport suggests a mechanism for tolerance/growth improvement by reducing the entry of aromatics. This is in line with a previous study in *P*. *putida* (9), which showed that deletion of an outer membrane porin PP_3350 in a wild- type strain decreased the lag phase in 20 g/L *p*-coumarate (∼123 mM) by >30 hours. In a recent study, Kusumawardhani *et al.* elucidated that several genes associated with porins and transport proteins were downregulated in an ALE-derived toluene-tolerant *P. putida* S12 (53). Besides the machinery associated with molecule uptake, efflux pumps have also been shown to contribute to the tolerance towards aromatic compounds (9, 49, 53). It is commonly known that aromatic compounds can disrupt cell membrane integrity due to their lipophilic (or partially lipophilic) nature (54, 55) and are also suggested to exert toxic effects intracellularly through different modes of actions (55, 56). Therefore, this mechanism of tolerance against aromatic compounds may result from their toxic effects in the periplasm or cytoplasm. In nature, the aromatic transport systems can be important for nutrient uptake, but in concentrations relevant to applications, their role becomes less important since aromatic acids in their protonated form can diffuse down the concentration gradient across cell membranes (35, 56). This may also explain our observation that wild-type ADP1 showed improved growth on ferulate at a higher pH, which can promote deprotonation and decrease the proportion of permeable aromatic acids.

The mutation in *ACIAD0482* was surprisingly found to be advantageous in ferulate tolerance. The product of *ACIAD0482* has not been reported in *A. baylyi* but has homology to LpsB of *Acinetobacter baumannii* with >80% identity. LpsB was reported to be a glycosyltransferase of the lipopolysaccharide (LPS) core (57). The 12 bp deletion in *ACIAD0482* would lead to the deletion of four amino acids from position 166 to 169 in the protein sequence. However, the effect of the deletion on the protein function is yet to be explored. Interestingly, another gene that is associated with lipooligosaccharide (LOS) was found to be mutated in ASA502 and ASA503: a single nucleotide deletion in *ACIAD0602*, which may lead to loss of protein function. *ACIAD0602* encodes a putative glycosyltransferase sharing >80% identity with GtrOC4 in *A*. *baumannii*, and GtrOC4 was proposed to be involved in the synthesis of the outer core of LOS (40). LPS/LOS is known to provide a barrier protecting gram-negative bacteria from hydrophobic substances (58, 59), but the mechanism of the tolerance improvement by the glycosyltransferase mutation remains unclear.

The IS*1236* element played an important role in the mutation development of the evolution experiment presented here. In addition to its insertion in *hcaE* in ASA500/ASA501, IS*1236* was found to be inserted in two non-coding regions in the strain ASA502 and ASA503. Interestingly, the gene duplications observed in ASA500 and ASA503 were found to be related to IS*1236*; the duplicated region was flanked by IS*1236*, but the genomic context of the original copy of the region seemed not to change. We speculated that the duplication resulted from the formation of a new composite transposon by IS*1236* flanking the duplicated region, followed by integration of the composite transposon into one of the IS*1236* sites. Although the duplications in the two strains originated from independent evolution events, and most of the duplicated genes were shared by the two strains, the roles of the duplications are not obvious. Because the strain ASA502, which was from the same population and shared many common mutations with ASA503, only showed a slight difference from ASA503 in the growth on ferulate but did not carry the duplication. *A. baylyi* ADP1 contains 6 copies of a single type of IS element, IS*1236*, five of which are identical (60). In a previous evolution study, it was reported that IS*1236* was responsible for 41% of mutations in ADP1 after propagation in rich nutrient broth for 1000 generations (61). Although IS elements may play a role in fitness improvement during evolution, they can also contribute to undesired genetic instability in engineered strains (42). This prompted us to use the transposon-free *A. baylyi* ADP1 (42), ISx, as the background strain for reverse engineering, though it seemed to have a decreased tolerance towards ferulate compared to wild-type ADP1.

From the point of view of rational engineering, it is desirable to find the “minimal set” of mutations resulting in significant improvement of a phenotype. Here, by employing the RAMSES methodology, we were rapidly able to identify and reintroduce two key mutations (the *hcaE* and *hcaK* mutations) that alone significantly improved the tolerance of *A. baylyi* ADP1 towards ferulate. Such a method would be particularly useful when screening a large number of mutations (and their combinations), in contrast to individual construction of knock-in cassettes. The high capability of screening would also make it possible to expand the subset of mutations to be tested beyond the mutations that are either intuitive or convergent between ALE replicates, increasing the potential to discover novel mechanisms behind the improved phenotypes. Here, for example, the *ACIAD0482* mutation, which may not be considered as beneficial intuitively, was found to be advantageous. In addition, the RAMSES approach can be easily automated with the use of a liquid handling robot, owing to the possibility of transforming *A. baylyi* directly in liquid culture.

## 4. Conclusion

We exploited the natural competence and high recombination efficiency of *A. baylyi* ADP1 in developing a simple and rapid method for screening, identifying, and reverse-engineering advantageous mutations that arose during ALE. The method was applied on strains that were evolved for high ferulate tolerance and then subjected to whole-genome sequencing. Among numerous mutations, we were able to determine that mutations in *hcaE* and *hcaK* played a major role in the improved tolerance. By simply introducing the combination of these two mutations in a parent strain, the high tolerance against ferulate could be restored. This study highlights the potential of applying the naturally competent *A. baylyi* ADP1 for evolution studies and strain development and facilitates the construction of more robust cell factories for aromatic substrate valorization.

## 5. Materials and methods

### 2.1. Strains and media

Wild-type *Acinetobacter baylyi* ADP1 (DSM 24193, DSMZ, Germany) was used as a starting strain for ALE, designated as WT ADP1. The transposon-free *A. baylyi* ADP1 (42) (a kind gift from Barrick lab), designated as ISx in this study, was used as the parent strain for reverse engineering. *E. coli* XL1-Blue (Stratagene, USA) was used as the host in cloning steps. All the strains used in this study are listed in Table S1.

Mineral salts medium (MSM) was used for ALE, growth study, and reverse engineering. The carbon sources, including ferulate, vanillate, *p*-coumarate, casamino acids, and acetate were added when appropriate. The composition of MSM was 3.88 g/l K_2_HPO_4_, 1.63 g/l NaH_2_PO_4_, 2.00 g/l (NH_4_)_2_SO_4_, 0.1 g/l MgCl_2_·6H_2_O, 10 mg/l Ethylenediaminetetraacetic acid (EDTA), 2 mg/l ZnSO_4_·7H_2_O, 1 mg/l CaCl_2_·2H_2_O, 5 mg/l FeSO_4_·7H_2_O, 0.2 mg/l Na_2_MoO_4_·2H_2_O, 0.2 mg/l CuSO_4_·5H_2_O, 0.4 mg/l CoCl_2_·6H_2_O, 1 mg/l MnCl_2_·2H_2_O. The stock solutions of ferulate, vanillate, and coumarate were prepared with a concentration of 200 mM; briefly, the proper amount of aromatic acid (all purchased from Sigma, USA) was added in deionized water and dissolved by slowly adding NaOH while stirring. The final pH of the stock solutions was 8.2∼8.3. The stock solutions were further sterilized by filtration with sterile filters (pore size 0.2 μm, Whatman). The stock solutions were freshly prepared before each experiment. *E. coli* strains were maintained on modified LB medium (10 g/L tryptone, 5 g/L yeast extract, 1 g/L NaCl) supplemented with 1% glucose. For solid medium, 15 g/L agar was added. Spectinomycin (50 µg/ml) was added when appropriate.

### 2.2. Adaptive laboratory evolution of *Acinetobacter baylyi* ADP1

Two parallel evolutions with ferulate as a sole carbon source, designated as G1 and G2 evolution lines here, have been described previously (23). Here, two additional parallel evolutions were carried out to improve the tolerance on ferulate, designated as T1 and T2 evolution lines, in which acetate and casamino acids were supplemented in addition to ferulate. The ALE cultivation was performed in Erlenmeyer flasks (100 ml) containing 10 ml medium at 30 °C and 300 rpm. Wild-type ADP1 was first plated on solid MSM, and 25 mM ferulate, 10 mM acetate, and 0.2% (W/V) casamino acids were supplemented. The single colony from the plate was pre-cultivated in MSM supplemented with 55 mM ferulate, 10 mM acetate, and 0.2% (W/V) casamino acids. The pre-culture was transferred to two Erlenmeyer flasks containing the same medium, resulting in the two parallel evolution populations. The cells were transferred to fresh media before reaching the stationary phase daily. The optical density at 600 nm (OD) was measured before each transfer. The amount of inoculum for each transfer was adjusted so that the initial OD after each transfer was between 0.03 and 0.1. The cells were cryopreserved at − 80 °C every two transfers. The concentration of ferulate was gradually increased during the evolution. Individual isolates were streak purified twice on LB-agar plates from the end population of each evolution line.

The number of generations (n) per flask was calculated with the following equation:

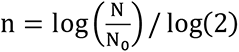

where N is the final OD_600_ of the culture and N_0_ is the initial OD_600_.

### 2.3. Phenotype characterization

The growth of different strains on different aromatic substrates was tested by cultivations in 96- well plates (Greiner Bio-One™ CellStar™ μClear™). The cells were pre-cultivated in MSM supplemented with 5 mM aromatic substrate (ferulate/vanillate/ *p*-coumarate) at which both the evolved strains and the reference strains can grow. After overnight cultivation, the cultures were inoculated (initial OD 0.05) into the media supplemented with the corresponding aromatic substrate at higher concentrations (as indicated in the result section). For strains from the evolution line T2, appropriate amounts of acetate and casamino acids were added when needed. The culture (200 µl) was transferred to the 96-well plate and incubated in Spark multimode microplate reader (Tecan, Switzerland) at 30 °C. Double orbital shaking was performed for 5 min twice an hour with an amplitude of 6 mm and a frequency of 54 rpm. OD was measured every hour. The cultivations were performed in duplicate. To study the effect of increased pH on cell growth, the pH of the media was adjusted by adding concentrated NaOH. The media was further sterilized by filtration. The cultivation was performed with the same procedure as mentioned above. For pre-cultivation, the media without pH adjustment was used.

### 2.4. Whole-genome sequencing of the evolved strains

Approximately 1 µg of genomic DNA from each strain was isolated using the Nucleospin gDNA cleanup kit (Macherey-Nigel), then fragmented by sonication to an average size of 300–500 bp. End repair, A-tailing, and adapter ligation reactions were performed on the fragmented DNA using the NEBNext Ultra II kit (New England Biolabs). Illumina paired-end sequencing was performed on a NextSeq500 device at the Georgia Genomics Facility (University of Georgia).

The sequences were analyzed using both Geneious prime version 8.1 with default settings (62) and the Breseq (version 0.35.4) computational pipeline (63). Version 2.4.1 of bowtie2 and version 4.0.0 of R were used in the pipeline. The consensus mode was used with the default consensus frequency cutoff of 0.8 and polymorphism frequency cutoff of 0.2. The raw reads from the five sequenced strains (ASA500, ASA501, ASA502, ASA503, and WT ADP1) were mapped to the reference genome of *A. baylyi* ADP1 (GenBank: CR543861).

### 2.5. Initial screening of advantageous mutations

The mutated alleles were first PCR-amplified from the evolved strains with Phusion high-fidelity DNA polymerase (Thermo Scientific, Finland), using the primers listed in Table S2. The amplified DNA fragments contained at least 500 bp of homology on each side of the mutated region. The PCR products were then loaded onto the agarose gel for electrophoresis. To avoid cross-contamination between the PCR products of the different mutated alleles, it is important to leave one well empty between the samples and not to overload the PCR products. The amplified DNA was purified with GeneJET Gel Extraction Kit (Thermo Scientific) and eluted with pre-warmed water. The concentrations of the purified PCR products ranged from 30 ng/µl to 100 ng/ µl. For natural transformation, ISx was first pre-cultivated in LB medium supplemented with 0.4% glucose. When the cells were in early exponential phase, 20 µl of the purified DNA was directly added to 180 µl of the culture, and then the mixture was incubated in 14 ml cultivation tube at 30 °C and 300 rpm for 3∼4 hours. The cells treated with water, an unmutated allele (gene entry: *ACIAD3383*) amplified from the evolved strain, and without any treatment were used as the controls. To adapt the cells to the medium used for the downstream process, 5 ml of MSM supplemented with 5 mM ferulate was added to the tube, and the culture was incubated overnight. After the incubation, 10 µl of the cells were transferred to different wells of a 96-well plate (Greiner Bio-One™ CellStar™ μClear™) containing 140 µl of MSM with elevated ferulate concentrations (20 mM – 80 mM). The plate was incubated in Spark multimode microplate reader (Tecan, Switzerland) at 30 °C, and the OD was measured every hour.

### 2.6. Reverse engineering of key mutations

The selected mutated alleles were PCR-amplified from the evolved strains using the primers listed in Table S2, and gel-extracted. For transformation, the background strain, ISx, was first streaked on LB-agar, and the plate was incubated at 30 °C overnight. The purified DNA (0.5 µl) was added onto single colonies and mixed well by pipetting up and down. After overnight incubation at 30 °C, the colony treated with the DNA was scraped and suspended in MSM supplemented 5 mM of the corresponding aromatic substrate (ferulate or vanillate). As the control, a colony without DNA treatment was subjected to the same process. The suspension was further incubated at 30 °C and 300 rpm for 0.5-10 h. After incubation, the suspension was used to inoculate 200 µl of MSM supplemented with elevated concentrations of the aromatic substrate (ferulate or vanillate) in different wells of the 96-well plate. The plate was incubated in Spark multimode microplate reader (Tecan, Switzerland) at 30 °C, and the OD was monitored every hour. If the cells treated with the mutated allele showed improved growth over the control at the elevated aromatic concentration, 5 µl of the cells were taken from the well and used to inoculate 5 ml of MSM containing the same (or higher) concentration of the corresponding aromatic substrate for further mutant enrichment. The culture was further streaked on LB-agar. The clones carrying the mutated allele were identified by picking single colonies for PCR analysis or Sanger sequencing.

### 2.7. Genetic engineering

ASA509 was constructed by transforming ASA504 with a linear integration cassette containing the spectinomycin resistance gene flanked by the sequences homologous to the sequences surrounding the *vanKP* region. The cassette was constructed by overlap extension PCR with the left flanking sequence (amplified with primers P1-F and P2-R, Table S2), the spectinomycin resistance gene (amplified with primers spec-F and spec-R), and the right flanking sequence (amplified with primers P3-F and P4-R). The linear cassette was later cloned to a previously described plasmid (18), and the left flanking sequence was replaced with another flanking sequence amplified with primers P5-F and P6-R. The resulting plasmid (non-replicating plasmid in ADP1) was used to transform ASA504 to obtain ASA510.

### 2.8. Analysis of substrate consumption

The concentrations of ferulate, vanillate, and acetate were analyzed using Agilent Technology 1100 Series HPLC (UV/VIS system) equipped with G1313A autosampler, G1322A degasser, G1311A pump, and G1315A DAD. Rezex RFQ-Fast Acid H+ (8%) (Phenomenex) was used as the column and placed at 80 °C. Sulfuric acid (0.005 N) was used as the eluent with a pumping rate of 0.8 ml/min.

## Data availability

Next-generation sequencing data generated in this study are available in NCBI Sequence Read Archive (SRA) BioProject accession number: PRJNA761218.

## Acknowledgements

Funding: The research work was supported by Academy of Finland (grants no. 310188, 334822), Novo Nordisk Foundation (grant no. NNF21OC0067758) and U.S. Department of Agriculture (grant no. 2018-67009-27926).

## Author contribution

Author contribution: JL, SS, and VS designed the study. JL and EM carried out the research work. JL, EM, and SB analyzed the data. SS, EN, and VS supervised the study. All authors participated in writing the manuscript.

## References

1. Dragosits M, Mattanovich D. 2013. Adaptive laboratory evolution – principles and applications for biotechnology. Microb Cell Fact 12:64.

2. Sandberg TE, Salazar MJ, Weng LL, Palsson BO, Feist AM. 2019. The emergence of adaptive laboratory evolution as an efficient tool for biological discovery and industrial biotechnology. Metab Eng 56:1–16.

3. Fletcher E, Feizi A, Bisschops MMM, Hallström BM, Khoomrung S, Siewers V, Nielsen J. 2017. Evolutionary engineering reveals divergent paths when yeast is adapted to different acidic environments. Metab Eng 39:19–28.

4. Kildegaard KR, Hallström BM, Blicher TH, Sonnenschein N, Jensen NB, Sherstyk S, Harrison SJ, Maury J, Herrgård MJ, Juncker AS, Forster J, Nielsen J, Borodina I. 2014. Evolution reveals a glutathione-dependent mechanism of 3-hydroxypropionic acid tolerance. Metab Eng 26:57–66.

5. Mundhada H, Seoane JM, Schneider K, Koza A, Christensen HB, Klein T, Phaneuf P V., Herrgard M, Feist AM, Nielsen AT. 2017. Increased production of L-serine in Escherichia coli through Adaptive Laboratory Evolution. Metab Eng 39:141–150.

6. Almario MP, Reyes LH, Kao KC. 2013. Evolutionary engineering of Saccharomyces cerevisiae for enhanced tolerance to hydrolysates of lignocellulosic biomass. Biotechnol Bioeng 110:2616– 2623.

7. Lim HG, Fong B, Alarcon G, Magurudeniya HD, Eng T, Szubin R, Olson CA, Palsson BO, Gladden JM, Simmons BA, Mukhopadhyay A, Singer SW, Feist AM. 2020. Generation of ionic liquid tolerant Pseudomonas putida KT2440 strains via adaptive laboratory evolution. Green Chem 22:5677–5690.

8. Phaneuf P V., Yurkovich JT, Heckmann D, Wu M, Sandberg TE, King ZA, Tan J, Palsson BO, Feist AM. 2020. Causal mutations from adaptive laboratory evolution are outlined by multiple scales of genome annotations and condition-specificity. BMC Genomics 21:514.

9. Mohamed ET, Werner AZ, Salvachúa D, Singer CA, Szostkiewicz K, Rafael Jiménez-Díaz M, Eng T, Radi MS, Simmons BA, Mukhopadhyay A, Herrgård MJ, Singer SW, Beckham GT, Feist AM. 2020. Adaptive laboratory evolution of Pseudomonas putida KT2440 improves p-coumaric and ferulic acid catabolism and tolerance. Metab Eng Commun 11:e00143.

10. Lee D-H, Palsson BØ. 2010. Adaptive Evolution of Escherichia coli K-12 MG1655 during Growth on a Nonnative Carbon Source, l-1,2-Propanediol. Appl Environ Microbiol 76:4158–4168.

11. Atsumi S, Wu T, Machado IMP, Huang W, Chen P, Pellegrini M, Liao JC. 2010. Evolution, genomic analysis, and reconstruction of isobutanol tolerance in Escherichia coli. Mol Syst Biol 6:449.

12. Neidle EL, Ornston LN. 1986. Cloning and expression of Acinetobacter calcoaceticus catechol 1,2- dioxygenase structural gene catA in Escherichia coli. J Bacteriol 168:815–820.

13. Santala S, Santala V. 2021. Acinetobacter baylyi ADP1—naturally competent for synthetic biology. Essays Biochem.

14. de Berardinis V, Vallenet D, Castelli V, Besnard M, Pinet A, Cruaud C, Samair S, Lechaplais C, Gyapay G, Richez C, Durot M, Kreimeyer A, Le Fèvre F, Schächter V, Pezo V, Döring V, Scarpelli C, Médigue C, Cohen GN, Marlière P, Salanoubat M, Weissenbach J. 2008. A complete collection of single-gene deletion mutants of Acinetobacter baylyi ADP1. Mol Syst Biol 4:174.

15. Santala V, Karp M, Santala S. 2016. Bioluminescence-based system for rapid detection of natural transformation. FEMS Microbiol Lett 363:fnw125.

16. Jiang X, Palazzotto E, Wybraniec E, Munro LJ, Zhang H, Kell DB, Weber T, Lee SY. 2020. Automating Cloning by Natural Transformation. ACS Synth Biol 9:3228–3235.

17. Tumen-Velasquez M, Johnson CW, Ahmed A, Dominick G, Fulk EM, Khanna P, Lee SA, Schmidt AL, Linger JG, Eiteman MA, Beckham GT, Neidle EL. 2018. Accelerating pathway evolution by increasing the gene dosage of chromosomal segments. Proc Natl Acad Sci 115:7105–7110.

18. Santala S, Efimova E, Kivinen V, Larjo A, Aho T, Karp M, Santala V. 2011. Improved Triacylglycerol Production in Acinetobacter baylyi ADP1 by Metabolic Engineering. Microb Cell Fact 10:36.

19. Lehtinen T, Efimova E, Santala S, Santala V. 2018. Improved fatty aldehyde and wax ester production by overexpression of fatty acyl-CoA reductases. Microb Cell Fact 17:19.

20. 20. Salmela M, Lehtinen T, Efimova E, Santala S, Santala V. 2019. Alkane and wax ester production from lignin related aromatic compounds. Biotechnol Bioeng bit.27005.

21. Luo J, Efimova E, Losoi P, Santala V, Santala S. 2020. Wax ester production in nitrogen-rich conditions by metabolically engineered Acinetobacter baylyi ADP1. Metab Eng Commun 10:e00128.

22. Santala S, Santala V, Liu N, Stephanopoulos G. 2021. Partitioning metabolism between growth and product synthesis for coordinated production of wax esters in Acinetobacter baylyi ADP1. Biotechnol Bioeng.

23. Luo J, Lehtinen T, Efimova E, Santala V, Santala S. 2019. Synthetic metabolic pathway for the production of 1-alkenes from lignin-derived molecules. Microb Cell Fact 18:48.

24. Arvay E, Biggs BW, Guerrero L, Jiang V, Tyo K. 2021. Engineering Acinetobacter baylyi ADP1 for mevalonate production from lignin-derived aromatic compounds. Metab Eng Commun 13:e00173.

25. Vardon DR, Franden MA, Johnson CW, Karp EM, Guarnieri MT, Linger JG, Salm MJ, Strathmann TJ, Beckham GT. 2015. Adipic acid production from lignin. Energy Environ Sci 8:617–628.

26. Salvachúa D, Karp EM, Nimlos CT, Vardon DR, Beckham GT. 2015. Towards lignin consolidated bioprocessing: simultaneous lignin depolymerization and product generation by bacteria. Green Chem 17:4951–4967.

27. 27. Linger JG, Vardon DR, Guarnieri MT, Karp EM, Hunsinger GB, Franden MA, Johnson CW, Chupka G, Strathmann TJ, Pienkos PT, Beckham GT. 2014. Lignin valorization through integrated biological funneling and chemical catalysis. Proc Natl Acad Sci.

28. Abdelaziz OY, Brink DP, Prothmann J, Ravi K, Sun M, García-Hidalgo J, Sandahl M, Hulteberg CP, Turner C, Lidén G, Gorwa-Grauslund MF. 2016. Biological valorization of low molecular weight lignin. Biotechnol Adv 34:1318–1346.

29. Beckham GT, Johnson CW, Karp EM, Salvachúa D, Vardon DR. 2016. Opportunities and challenges in biological lignin valorization. Curr Opin Biotechnol 42:40–53.

30. Harwood CS, Parales RE. 1996. The β-ketoadipate Pathway and the Biology of Self-identity. Annu Rev Microbiol 50:553–590.

31. Fischer R, Bleichrodt FS, Gerischer UC. 2008. Aromatic degradative pathways in Acinetobacter baylyi underlie carbon catabolite repression. Microbiology 154:3095–3103.

32. 32. Seaton SC, Neidle EL. 2018. Chapter 10. Using Aerobic Pathways for Aromatic Compound Degradation to Engineer Lignin Metabolism, p. 252–289. In.

33. Pardo I, Jha RK, Bermel RE, Bratti F, Gaddis M, McIntyre E, Michener W, Neidle EL, Dale T, Beckham GT, Johnson CW. 2020. Gene amplification, laboratory evolution, and biosensor screening reveal MucK as a terephthalic acid transporter in Acinetobacter baylyi ADP1. Metab Eng 62:260–274.

34. Smith MA, Weaver VB, Young DM, Ornston LN. 2003. Genes for Chlorogenate and Hydroxycinnamate Catabolism (hca) Are Linked to Functionally Related Genes in the dca-pca-qui- pob-hca Chromosomal Cluster of Acinetobacter sp. Strain ADP1. Appl Environ Microbiol 69:524–532.

35. Nichols NN, Harwood CS. 1997. PcaK, a high-affinity permease for the aromatic compounds 4- hydroxybenzoate and protocatechuate from Pseudomonas putida. J Bacteriol 179:5056–5061.

36. D’Argenio DA, Segura A, Coco WM, Bünz P V., Ornston LN. 1999. The Physiological Contribution ofAcinetobacter PcaK, a Transport System That Acts upon Protocatechuate, Can Be Masked by the Overlapping Specificity of VanK. J Bacteriol 181:3505–3515.

37. Parke D, Ornston LN. 2003. Hydroxycinnamate (hca) Catabolic Genes from Acinetobacter sp. Strain ADP1 Are Repressed by HcaR and Are Induced by Hydroxycinnamoyl-Coenzyme A Thioesters. Appl Environ Microbiol 69:5398–5409.

38. Morawski B, Segura A, Ornston LN. 2000. Substrate Range and Genetic Analysis of Acinetobacter Vanillate Demethylase. J Bacteriol 182:1383–1389.

39. Cerqueira GM, Kostoulias X, Khoo C, Aibinu I, Qu Y, Traven A, Peleg AY. 2014. A Global Virulence Regulator in Acinetobacter baumannii and Its Control of the Phenylacetic Acid Catabolic Pathway. J Infect Dis 210:46–55.

40. Kenyon JJ, Nigro SJ, Hall RM. 2014. Variation in the OC Locus of Acinetobacter baumannii Genomes Predicts Extensive Structural Diversity in the Lipooligosaccharide. PLoS One 9:e107833.

41. Gerischer U, D’Argenio DA, Ornston LN. 1996. IS 1236, a newly discovered member of the IS3 family, exhibits varied patterns of insertion into the Acinetobacter calcoaceticus chromosome. Microbiology 142:1825–1831.

42. Suárez GA, Renda BA, Dasgupta A, Barrick JE. 2017. Reduced Mutation Rate and Increased Transformability of Transposon-Free Acinetobacter baylyi ADP1-ISx. Appl Environ Microbiol 83.

43. Ragauskas AJ. 2006. The Path Forward for Biofuels and Biomaterials. Science (80-) 311:484–489.

44. Schutyser W, Renders T, Van den Bosch S, Koelewijn S-F, Beckham GT, Sels BF. 2018. Chemicals from lignin: an interplay of lignocellulose fractionation, depolymerisation, and upgrading. Chem Soc Rev 47:852–908.

45. Ragauskas AJ, Beckham GT, Biddy MJ, Chandra R, Chen F, Davis MF, Davison BH, Dixon RA, Gilna P, Keller M, Langan P, Naskar AK, Saddler JN, Tschaplinski TJ, Tuskan GA, Wyman CE. 2014. Lignin Valorization: Improving Lignin Processing in the Biorefinery. Science (80-) 344:1246843–1246843.

46. Biggs BW, Bedore SR, Arvay E, Huang S, Subramanian H, McIntyre EA, Duscent-Maitland C V., Neidle EL, Tyo KEJ. 2020. Development of a genetic toolset for the highly engineerable and metabolically versatile Acinetobacter baylyi ADP1. Nucleic Acids Res 48:5169–5182.

47. Kohlstedt M, Starck S, Barton N, Stolzenberger J, Selzer M, Mehlmann K, Schneider R, Pleissner D, Rinkel J, Dickschat JS, Venus J, B.J.H. van Duuren J, Wittmann C. 2018. From lignin to nylon: Cascaded chemical and biochemical conversion using metabolically engineered Pseudomonas putida. Metab Eng 47:279–293.

48. Salvachúa D, Johnson CW, Singer CA, Rohrer H, Peterson DJ, Black BA, Knapp A, Beckham GT. 2018. Bioprocess development for muconic acid production from aromatic compounds and lignin. Green Chem 20:5007–5019.

49. Calero P, Jensen SI, Bojanovič K, Lennen RM, Koza A, Nielsen AT. 2018. Genome-wide identification of tolerance mechanisms toward p -coumaric acid in Pseudomonas putida. Biotechnol Bioeng 115:762–774.

50. van Duuren JBJH, Wijte D, Karge B, Martins dos Santos VAP, Yang Y, Mars AE, Eggink G. 2012. pH- stat fed-batch process to enhance the production of cis, cis-muconate from benzoate by Pseudomonas putida KT2440-JD1. Biotechnol Prog 28:85–92.

51. Pernstich C, Senior L, MacInnes KA, Forsaith M, Curnow P. 2014. Expression, purification and reconstitution of the 4-hydroxybenzoate transporter PcaK from Acinetobacter sp. ADP1. Protein Expr Purif 101:68–75.

52. Kim Y, Joachimiak G, Bigelow L, Babnigg G, Joachimiak A. 2016. How Aromatic Compounds Block DNA Binding of HcaR Catabolite Regulator. J Biol Chem 291:13243–13256.

53. Kusumawardhani H, Furtwängler B, Blommestijn M, Kaltenytė A, van der Poel J, Kolk J, Hosseini R, de Winde JH. 2021. Adaptive Laboratory Evolution Restores Solvent Tolerance in Plasmid- Cured Pseudomonas putida S12: a Molecular Analysis. Appl Environ Microbiol 87:1–18.

54. Ramos JL, Duque E, Gallegos M-T, Godoy P, Ramos-González MI, Rojas A, Terán W, Segura A. 2002. Mechanisms of Solvent Tolerance in Gram-Negative Bacteria. Annu Rev Microbiol 56:743– 768.

55. Mills TY, Sandoval NR, Gill RT. 2009. Cellulosic hydrolysate toxicity and tolerance mechanisms in Escherichia coli. Biotechnol Biofuels 2:26.

56. Borges A, Ferreira C, Saavedra MJ, Simões M. 2013. Antibacterial Activity and Mode of Action of Ferulic and Gallic Acids Against Pathogenic Bacteria. Microb Drug Resist 19:256–265.

57. Luke NR, Sauberan SL, Russo TA, Beanan JM, Olson R, Loehfelm TW, Cox AD, St. Michael F, Vinogradov E V., Campagnari AA. 2010. Identification and Characterization of a Glycosyltransferase Involved in Acinetobacter baumannii Lipopolysaccharide Core Biosynthesis. Infect Immun 78:2017–2023.

58. Zhang G, Baidin V, Pahil KS, Moison E, Tomasek D, Ramadoss NS, Chatterjee AK, McNamara CW, Young TS, Schultz PG, Meredith TC, Kahne D. 2018. Cell-based screen for discovering lipopolysaccharide biogenesis inhibitors. Proc Natl Acad Sci 115:6834–6839.

59. May KL, Grabowicz M. 2018. The bacterial outer membrane is an evolving antibiotic barrier. Proc Natl Acad Sci 115:8852–8854.

60. Cuff LE, Elliott KT, Seaton SC, Ishaq MK, Laniohan NS, Karls AC, Neidle EL. 2012. Analysis of is1236-mediated gene amplification events in acinetobacter baylyi ADP1. J Bacteriol.

61. Renda BA, Dasgupta A, Leon D, Barrick JE. 2015. Genome instability mediates the loss of key traits by Acinetobacter baylyi ADP1 during laboratory evolution. J Bacteriol.

62. Kearse M, Moir R, Wilson A, Stones-Havas S, Cheung M, Sturrock S, Buxton S, Cooper A, Markowitz S, Duran C, Thierer T, Ashton B, Meintjes P, Drummond A. 2012. Geneious Basic: An integrated and extendable desktop software platform for the organization and analysis of sequence data. Bioinformatics.

63. 63. Deatherage DE, Barrick JE. 2014. Identification of Mutations in Laboratory-Evolved Microbes from Next-Generation Sequencing Data Using breseq, p. 165–188. *In*.

