## Supplemental materials for "Characterization of highly ferulate-tolerant *Acinetobacter baylyi* ADP1 isolates by a rapid reverse-engineering method"

Jin Luo^1^#, Emily A. McIntyre^2^, Stacy R. Bedore^2^, Ville Santala^1^, Ellen L. Neidle^2^, Suvi Santala^1^
^1^Faculty of Engineering and Natural Sciences, Hervanta campus, Tampere University, Korkeakoulunkatu 8, Tampere, 33720, Finland
^2^Department of Microbiology, University of Georgia, Athens, GA 30602-2605 USA

Emily A. McIntyre:
Stacy R. Bedore:
Ville Santala:
Ellen L. Neidle:
Suvi Santala:

**Supplemental Materials**

**Table S1.** Bacterial strains used in the study

| Name | Description | Source / reference |
| --- | --- | --- |
| *E. coli* XL1-Blue | Wild-type *E. coli* XL1-Blue | Stratagene, USA |
| WT ADP1 | Wild-type *A. baylyi* ADP1 | DSM 24193, DSMZ |
| ASA500 | isolate from G1 evolution line | (1) |
| ASA501 | isolate from G2 evolution line | (1) |
| ASA502 | isolate from T2 evolution line | this study |
| ASA503 | isolate from T2 evolution line | this study |
| ISx | Transposon-free *A. baylyi* ADP1 | (2) |
| ASA504 | reconstructed mutant containing *hcaE500*, descended from ISx, constructed by RAMSES with ferulate | this study |
| ASA505 | reconstructed mutant containing *vanK500*, descended from ISx, constructed by RAMSES with ferulate | this study |
| ASA506 | reconstructed mutant containing *hcaE500* and *vanK500*, descended from ASA504, constructed by RAMSES with vanillate | this study |
| ASA507 | reconstructed mutant containing *hcaE500* and *hcaK501*, descended from ASA504, constructed by RAMSES with ferulate | this study |
| ASA508 | reconstructed mutant containing *hcaE500* and *ACIAD0482_500*, descended from ASA504, constructed by RAMSES with ferulate | this study |
| ASA509 | ∆*vanKP*::spec^r^ mutant descended from ASA504, 732 bp region upstream of *vanK* was also deleted | this study |
| ASA510 | ∆*vanP*::spec^r^ mutant descended from ASA504 | this study |

**Table S2.** Primers used in the study

| Name | Sequence | Description |
| --- | --- | --- |
| pcaU-M-F | CATCAGGGCTGACTGCTGAA | forward primer used to amplify *pcaU500* |
| pcaU-M-R | CAATTTTGCCCGCACGGTAT | reverse primer used to amplify *pcaU500* |
| hcaE-M-F | GCCTTTGAGCTGAGCACCTA | forward primer used to amplify *hcaE500* |
| hcaE-M-R | AGTAACTCAGCGCCTTGGTC | reverse primer used to amplify *hcaE500* |
| hcaK-M-F | AAATCCATGCCAGCAGTCCA | forward primer used to amplify *hcaK501* |
| hcaK-M-R | ACGGCATTTGATTTTGCCCA | reverse primer used to amplify *hcaK501* |
| vanK-M-F | TTCCACGTAACGCCATTTGC | forward primer used to amplify *vanK500* |
| vanK-M-R | GATATGCGCCACCCAAGGTA | reverse primer used to amplify *vanK500* |
| ACIAD2867-M-F | AGTTTGGCTGAGTTGCCAGT | forward primer used to amplify *ACIAD2867_500* |
| ACIAD2867-M-R | GGCGTTCTGAACAATGCGAG | reverse primer used to amplify *ACIAD2867_500* |
| ACIAD2265-M-F | TGGCAACGTAACCAACCAGT | forward primer used to amplify *ACIAD2265_501* |
| ACIAD2265-M-R | ACATTGCTACAGCCAGCACT | reverse primer used to amplify *ACIAD2265_501* |
| ACIAD0482-M-F | TGGTCTTGGTTTACGCAGCA | forward primer used to amplify *ACIAD0482_500* |
| ACIAD0482-M-R | TCGTTGCCGTGCAACTATCT | reverse primer used to amplify *ACIAD0482_500* |
| P1-F | ATGGTACCCACACTGGATATGAAACAGC | forward primer used to amplify the left flanking sequence for *vanKP* knock-out, contains KpnI site |
| P2-R | ATACTCGAGATTAAATAAAATAGGATTGACTCTGC | reverse primer used to amplify the left flanking sequence for *vanKP* knock-out, contains XhoI site |
| P3-F | TATCCTAGGTCATTCAACCAACGAAACACG | forward primer used to amplify the right flanking sequence for *vanKP* and *vanP* knock-out, contains AvrII site |
| P4-R | ATTACGAATTCTTCATCCAAACGCAAAAGCC | reverse primer used to amplify the right flanking sequence for *vanKP* and *vanP* knock-out, contains EcoRI site |
| P5-F | TAGGTACCGCTGGACACCAAAAATTCTGA | forward primer used to amplify the left flanking sequence for *vanP* knock-out, contains KpnI site |
| P6-R | ATTACTCGAGTTAAGCAACTGTTTTTATAGAACTTC | reverse primer used to amplify the left flanking sequence for *vanP* knock-out, contains XhoI site |
| spec-F | GTCAATCCTATTTTATTTAATCTCGAGTATGCAGAAAGGAGAAGCTTACTAG | forward primer used to amplify the spectinomycin resistance gene, contains XhoI site |
| spec-R | CGTTGGTTGAATGACCTAGGATAGCTTAATGCGCCGCTACA | reverse primer used to amplify the spectinomycin resistance gene, contains AvrII site |

**Table S3.** Mutations present in the WT A. baylyi ADP1 used that differ from the NCBI deposited sequence: CR543861 (GeneBank entry). The mutation highlighted in grey is the only mutation that is not shared by the evolved isolates.

| **Gene locus ID (name) ^a^** | **Position ^a^** | **Description ^a^** | **Mutation type** | **DNA change** | **Protein effect** |
| --- | --- | --- | --- | --- | --- |
| N/A | 3357842 | Non-coding region between *ACIAD3437* and *ACIAD3440* | Substitution (transition) | G 🡪 A | N/A |
| *ACIAD2866* | 2803339-2805228 | Hemagglutinin/hemolysin‑related protein | Duplication | Repetition from the CDS position 1093 to 2982 | Repetition from the amino acid position 365 to 994 |
| *ACIAD2827* | 2765010 | Putative periplasmic binding protein of transport/transglycosylase | Substitution (transversion) | A 🡪 C | S353R  (AGT🡪CGT) |
| *ACIAD2584* (*lepA*) | 2542652 | GTP‑binding protein | Substitution (transition) | G 🡪 A | None |
| N/A | 2420279-2420280 | Non-coding region between *ACIAD2457* and *ACIAD2458* (*glnA*) | Insertion (tandem repeat) | (C)2 🡪 (C)3 | N/A |
| *ACIAD1796* | 1803939 | hypothetical protein | Substitution (transition) | C 🡪 T | None |
| *ACIAD1796* | 1803933 | hypothetical protein | Substitution (transition) | A 🡪 G | None |
| *ACIAD1796* | 1803930 | hypothetical protein | Substitution (transition) | T 🡪 C | None |
| *ACIAD1796* | 1803921 | hypothetical protein | Substitution (transition) | A 🡪 G | None |
| *ACIAD1796* | 1803915 | hypothetical protein | Substitution (transition) | G 🡪 A | None |
| *ACIAD1796* | 1803903 | hypothetical protein | Substitution (transition) | T 🡪 C | None |
| *ACIAD1796* | 1803900 | hypothetical protein | Substitution (transition) | C 🡪 T | None |
| *ACIAD1796* | 1803861 | hypothetical protein | Substitution (transition) | T 🡪 C | None |
| *ACIAD1796* | 1803858 | hypothetical protein | Substitution (transition) | C 🡪 T | None |
| *ACIAD1182* (*phrB*) | 1179108 | deoxyribodipyrimidine photolyase (photoreactivation) | Substitution (transition) | C 🡪 T | D388N  (GAC→AAC) |
| N/A | 178459 | Non-coding region between *ACIAD0177* (*znuA*) and *ACIAD0178* (atpI) | Deletion (tandem repeat) | (G)10 🡪 (G)9 | N/A |

^a^ Locus IDs, mutation positions, and descriptions were assigned according to the reference genome CR543861 (GeneBank entry).

**Table S4.** Details of the duplicated regions found in ASA500 and ASA503.

| **Strain** | **Coverage of genome** | **Coverage of the duplication region** | **Start position ^a^** | **End position ^a^** | **Length of the duplication region (bp)** | **Genes involved** |
| --- | --- | --- | --- | --- | --- | --- |
| ASA500 | 138.7 | 295.4 | 3380313 | 3408297 | 27985 | *ACIAD3457*-*ACIAD3481* |
| ASA503 | 115.4 | 238.6 | 3380938 | 3413938 | 33001 | *ACIAD3459*-*ACIAD3486* |

^a^ Positions assigned according the reference genome CR543861 (GeneBank entry)

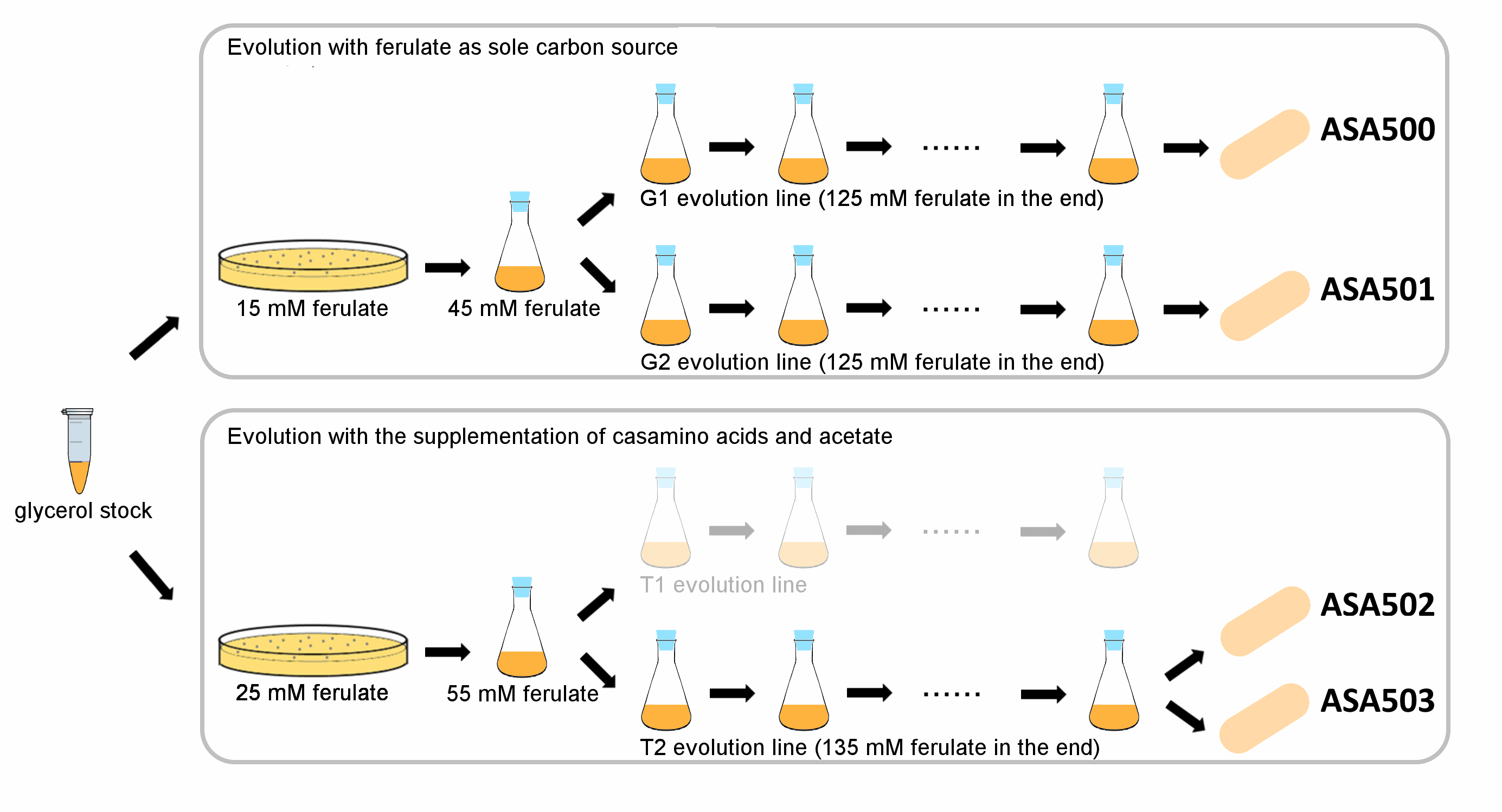
**Figure S1.** Outline of the ALE procedure. The experiments were initiated by streaking the cells to plates containing ferulate at lower concentrations, followed by pre-cultivation from single colonies with the starting ferulate concentrations. Four individual isolates from the endpoint populations streak purified, named, and used for subsequent studies.

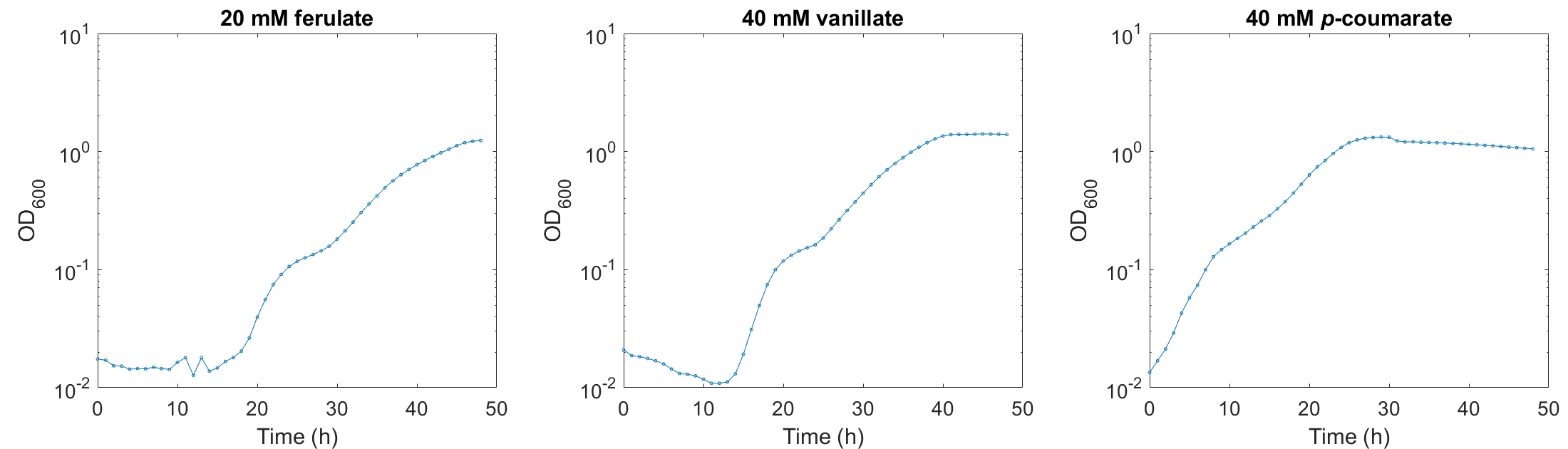

**Figure S2.** Typical examples showing the biphasic growth pattern of some *A. baylyi* ADP1 strains. A biphasic growth pattern was observed when ISx was grown in 20 mM ferulate, and when ASA504 was grown in 40 mM vanillate and 40 mM *p*-coumarate. All the strains were cultivated in mineral salts media with the corresponding aromatic compound as the sole carbon source. The y-axis is shown in log10 scale.

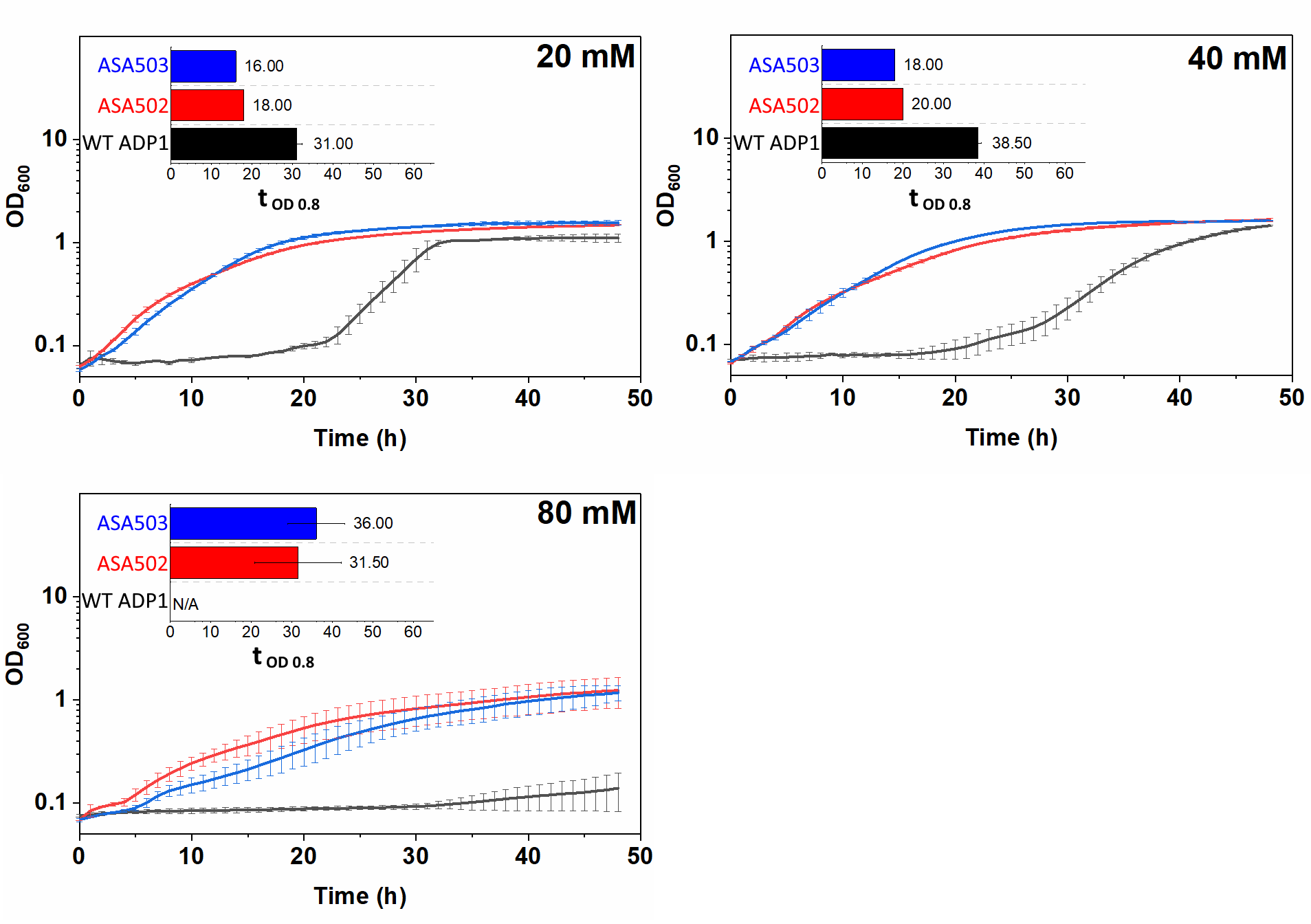
**Figure S3.** Growth of the strains ASA502, ASA503, and WT ADP1 at different ferulate concentrations (20 mM, 40 mM, and 80 mM). WT ADP1 was used as the control. All the strains were cultivated in mineral salts medium supplemented with ferulate as the sole carbon source. The time required for the cells to reach OD 0.8 was used as the indicator to evaluate the tolerance (t_OD 0.8_). The values were calculated from two replicates and the error bars indicate the standard deviations. The y-axis is shown in log10 scale.

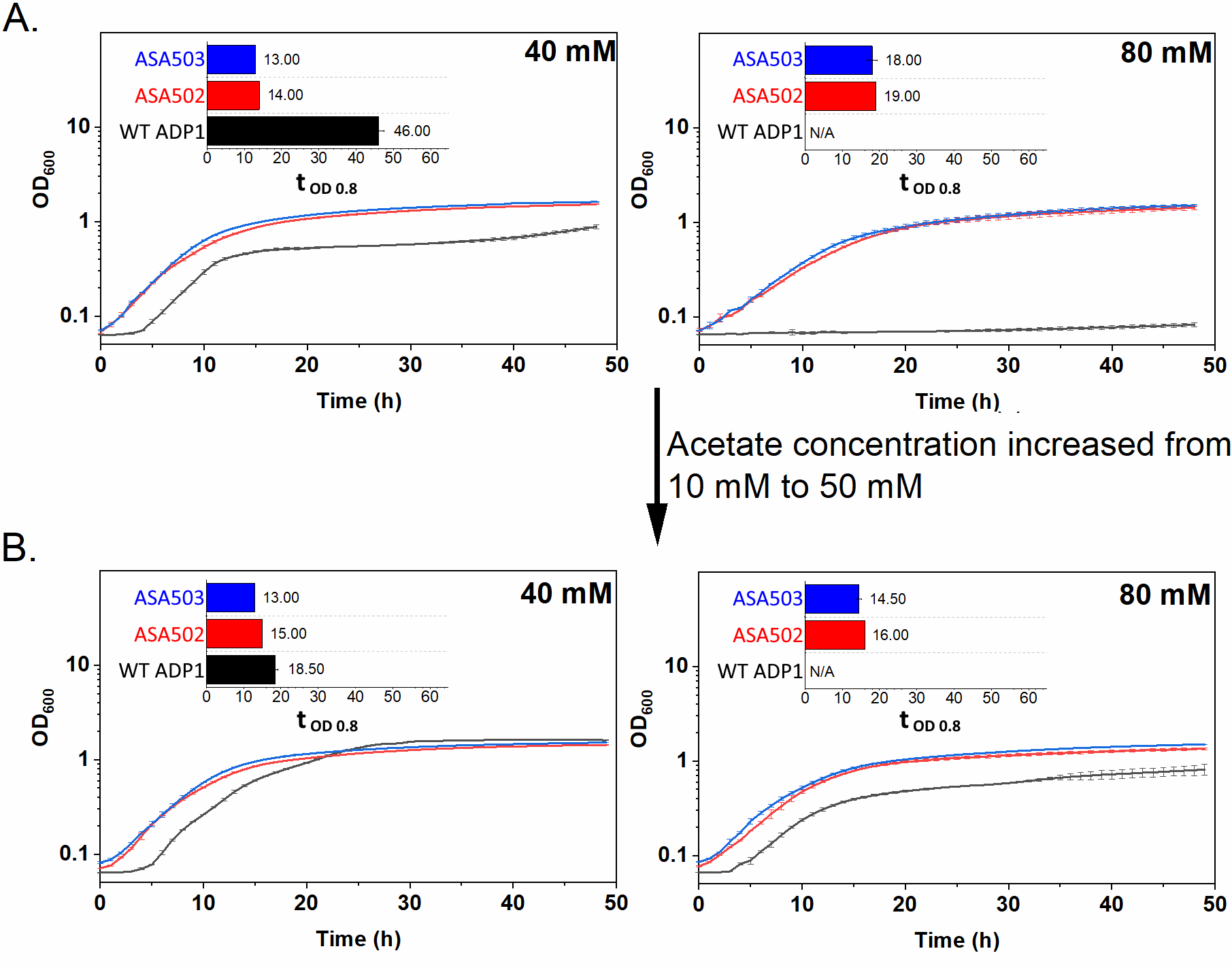
**Figure S4.** Growth of the strains ASA502, ASA503, and WT ADP1 at 40 mM and 80 mM of ferulate with the supplementation of casamino acids and acetate. All the strains were cultivated in mineral salts media supplemented with 0.2% (w/v) casamino acids and 10 mM (A) or 50 mM (B) acetate. The time required for the cells to reach OD of 0.8 was used as the indicator to evaluate the tolerance (t_OD 0.8_). The indicator was calculated only when both replicates reached OD 0.8 within 48 h. All the values were calculated from two replicates and the error bars indicate the standard deviations. The y-axis is shown in log10 scale, and the ticks for values from 0.1 to 10 are shown.

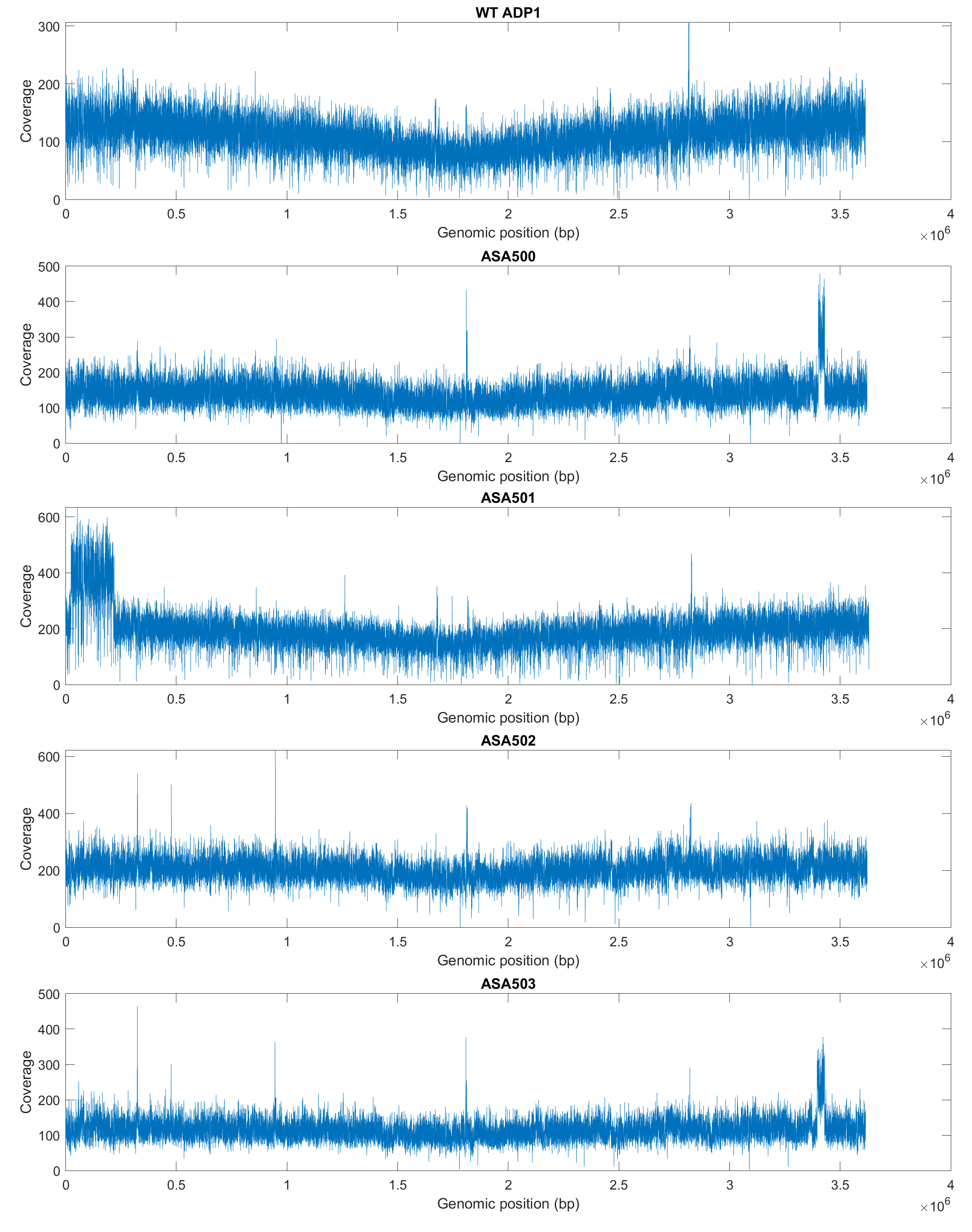
**Figure S5.** Sequencing coverages for the strains WT ADP1, ASA500, ASA501, ASA502, and ASA503.

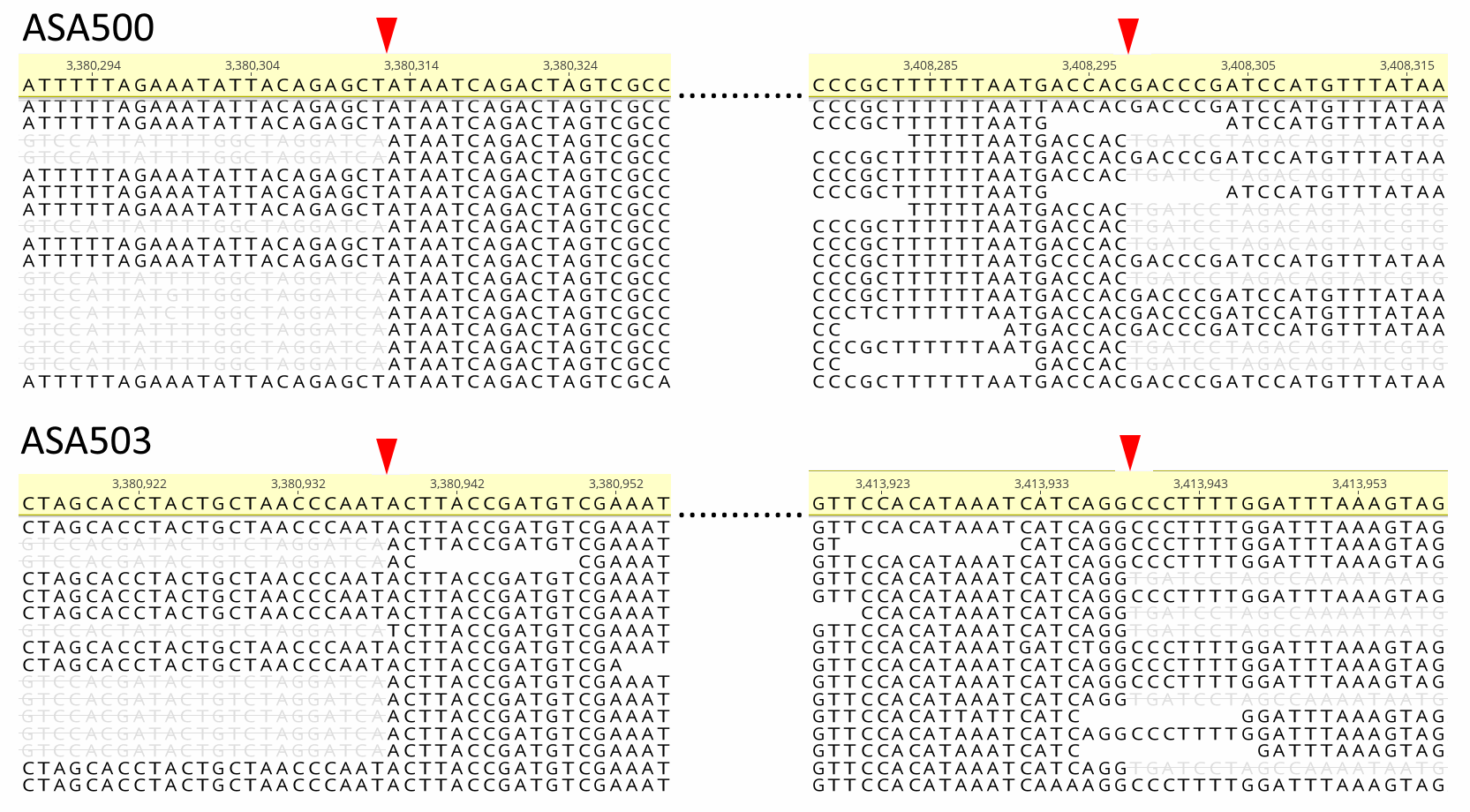
**Figure S6.** Mapping of the reads at the junctions of the duplicated regions in ASA500 and ASA503. The reads were mapped to the reference genome of *A. baylyi* ADP1 (GenBank: CR543861). The junction sites are indicated with red arrows, and the sequences of the IS element are indicated with strikethrough texts.

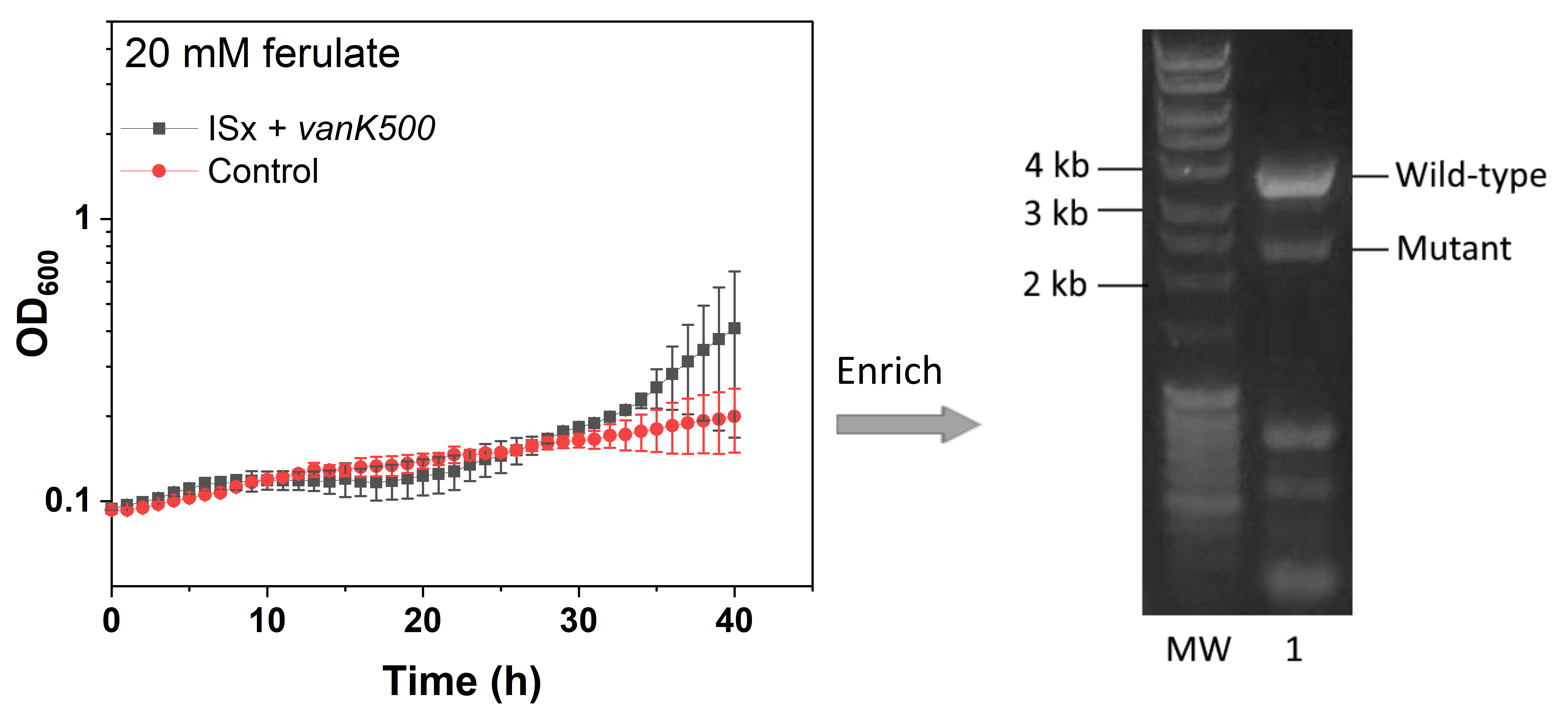
**Figure S7.** Reverse engineering of the mutated allele *vanK500* (loss of function by deletion, from ASA500) into ISx by RAMSES. Growth of ISx in 20 mM ferulate after being transformed with *vanK500* is shown. Control: ISx without transformation. After further enrichment, PCR was performed to amplify the *vanK* region from the genome extracted from the *vanK500* transformed population. Lane 1: the transformed population. The transformation was done with solid medium. The OD values were calculated from two replicates. The error bars indicate the standard deviations. The y-axis is shown in log10 scale, and the ticks for values from 0.1 to 5 are shown.

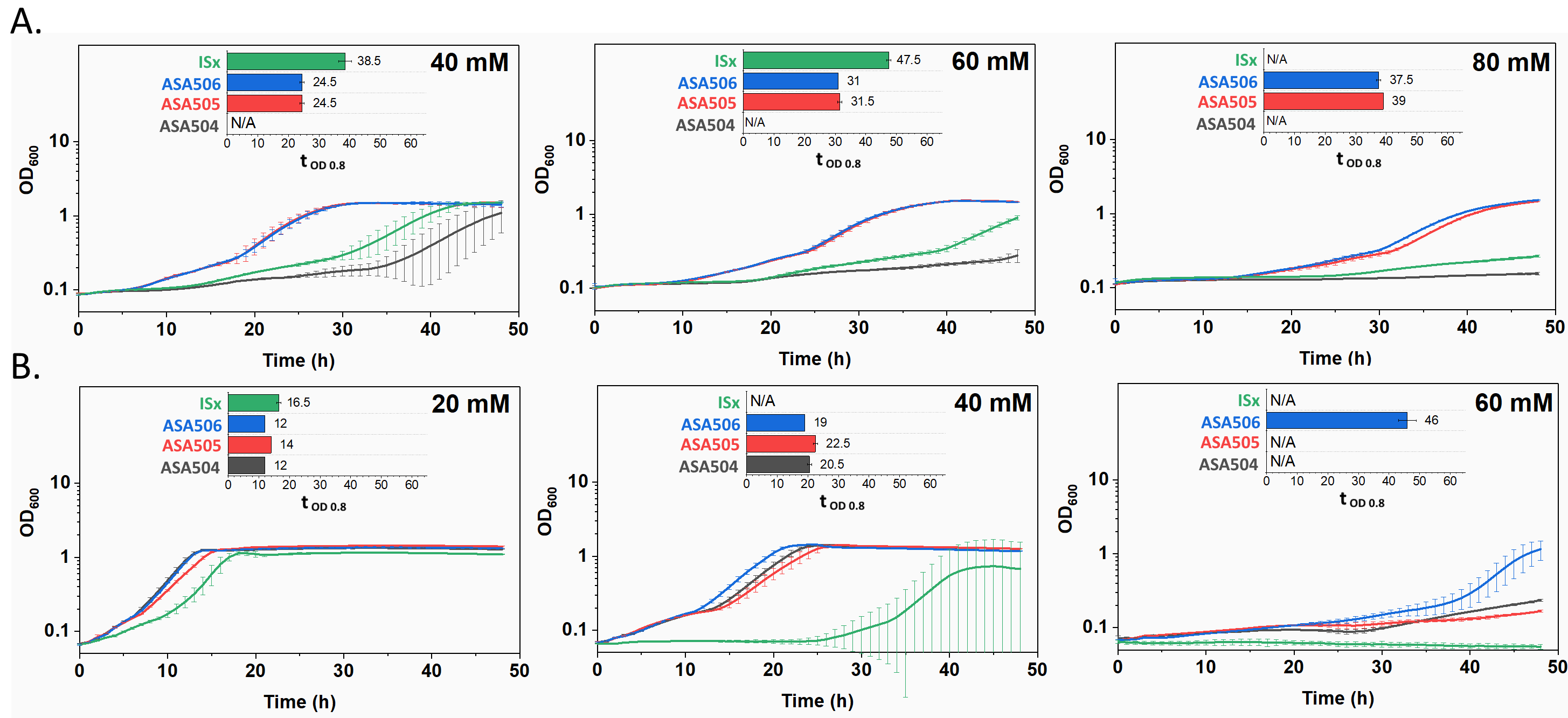
**Figure S8.** Growth comparison between ISx, ASA504 (reconstructed mutant *hcaE*), ASA505 (reconstructed mutant *vanK*), and ASA506 (reconstructed mutant *hcaE* and *vanK*) at different vanillate and *p*-courmarate concentrations. All the strains were cultivated in mineral salts media with vanillate (A) or *p*-courmarate (B) as the sole carbon source. Time spent for the cells to reach the OD of 0.8 was used as the indicator to evaluate the tolerance (t_OD 0.8_). The indicator is calculated only when both replicates reached OD 0.8 within 48 h. All the values were calculated from two replicates and the error bars indicate the standard deviations. The y-axis is shown in log10 scale.

**Supplemental Note**

**The effect of pH on growth on ferulate**

Compared to the evolved strains, WT ADP1showed a larger variance in aromatic tolerance between different independent experiments even though the same concentration of the aromatic compounds was supplied. On one hand, WT ADP1 could be more sensitive to the toxicity of aromatic compounds than the evolved strains; on the other hand, a slight difference in the growth condition might affect the toxicity to cells. Aromatic acid in its undissociated form can diffuse across the cell membrane (3). Therefore, the pH of the medium might be one of the factors that would alter the toxicity of the aromatic acid. Hence, we examined the effect of pH on cell growth on ferulate. Both the evolved strain ASA500 and WT ADP1 were cultivated with the supplementation of 40 mM and 80 mM ferulate under two pH conditions. In the first condition, the pH of the media was not adjusted, which resulted in a pH of 7.12 (media with 40 mM ferulate) and 7.22 (media with 80 mM ferulate). In the second condition, the pH of the media was increased, within the tolerable range, to 8.42 by adding NaOH. A recognizable difference in growth was observed for WT ADP1 between the two conditions (Figure S9). At 40 mM of ferulate, the increase of the pH reduced the time for WT ADP1 to reach the OD of 0.8 from 30.5 h to 21.5 h; at 80 mM, WT ADP1 was able to grow at the pH of 8.42 but not 7.22. In contrast, ASA500 did not show a large difference in growth at the two pH conditions (Figure S9); its growth seemed to be negatively affected at the higher pH.

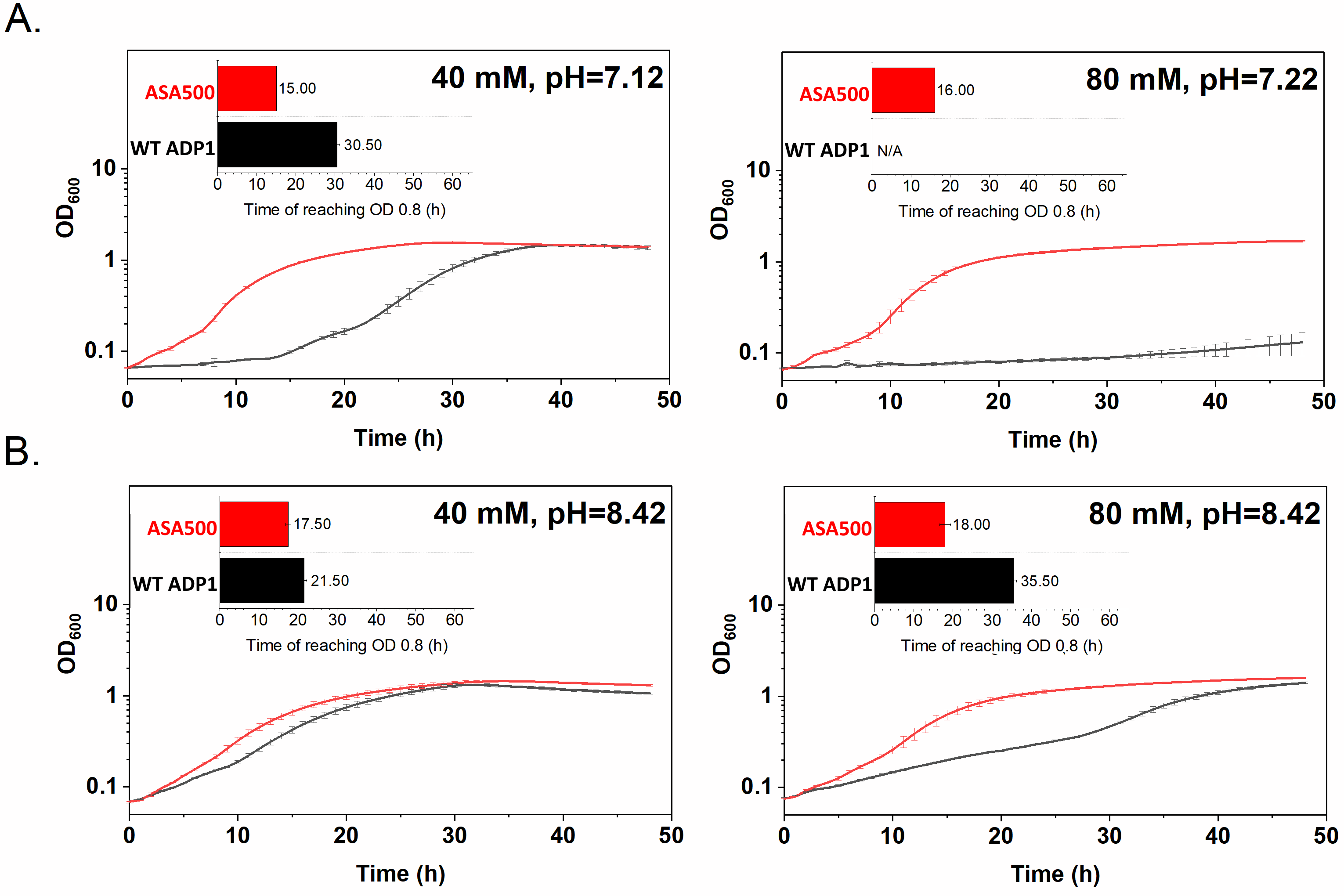

**Figure S9.** The effect of pH on the growth of cells on ferulate. Growth of the evolved strain ASA500 and WT ADP1 in mineral salts media supplemented with 40 mM and 80 mM of ferulate before pH adjustment (A) and after pH was adjusted to 8.42 (B). Time spent for the cells to reach the OD of 0.8 was used as the indicator to evaluate the tolerance (t_OD 0.8_). All the values were calculated from two replicates and the error bars indicate the standard deviations. The y-axis is shown in log10 scale.

**The *vanK* mutation has a different effect than deletion of either *vanKP* or *vanP***

The *vanK* mutation (1137 bp deletion, from ASA500) resulted from a deletion from the position 732 bp upstream of *vanK* to its CDS position 405, indicating that the promoter region of *vanK* might be affected. *vanP*, proposed to encode an outer membrane porin for aromatic uptake (4), is only 31 bp downstream of *vanK*, suggesting that it might be transcribed together with *vanK*. To inspect if deletion of both *vanK* and *vanP* or only *vanP* had the same effect as the *vanK* mutation to restore the tolerance of ASA504 (reconstructed mutant *hcaE*) to vanillate, two strains were constructed using ASA504 (reconstructed mutant *hcaE*) as the parental strain: ASA509, in which the 732 bp region upstream of *vanK*, *vanK* CDS and *vanP* CDS were replaced with the spectinomycin resistance gene, and ASA510, in which *vanP* CDS was replaced with the spectinomycin resistance gene (Figure S10A). Although both ASA509 and ASA510 showed improved tolerance towards vanillate compared to ASA504 (reconstructed mutant *hcaE*), they did not grow as well as ASA506 (reconstructed *hcaE* and *vanK* mutant) (Figure S10B). The result indicates that the *vanK* mutation (large deletion, from ASA500) had a different effect than deletion of both *vanK* and *vanP* or only *vanP*. But it cannot be excluded that the change of the genomic context or the introduction of the antibiotic marker may have an effect on the tolerance.

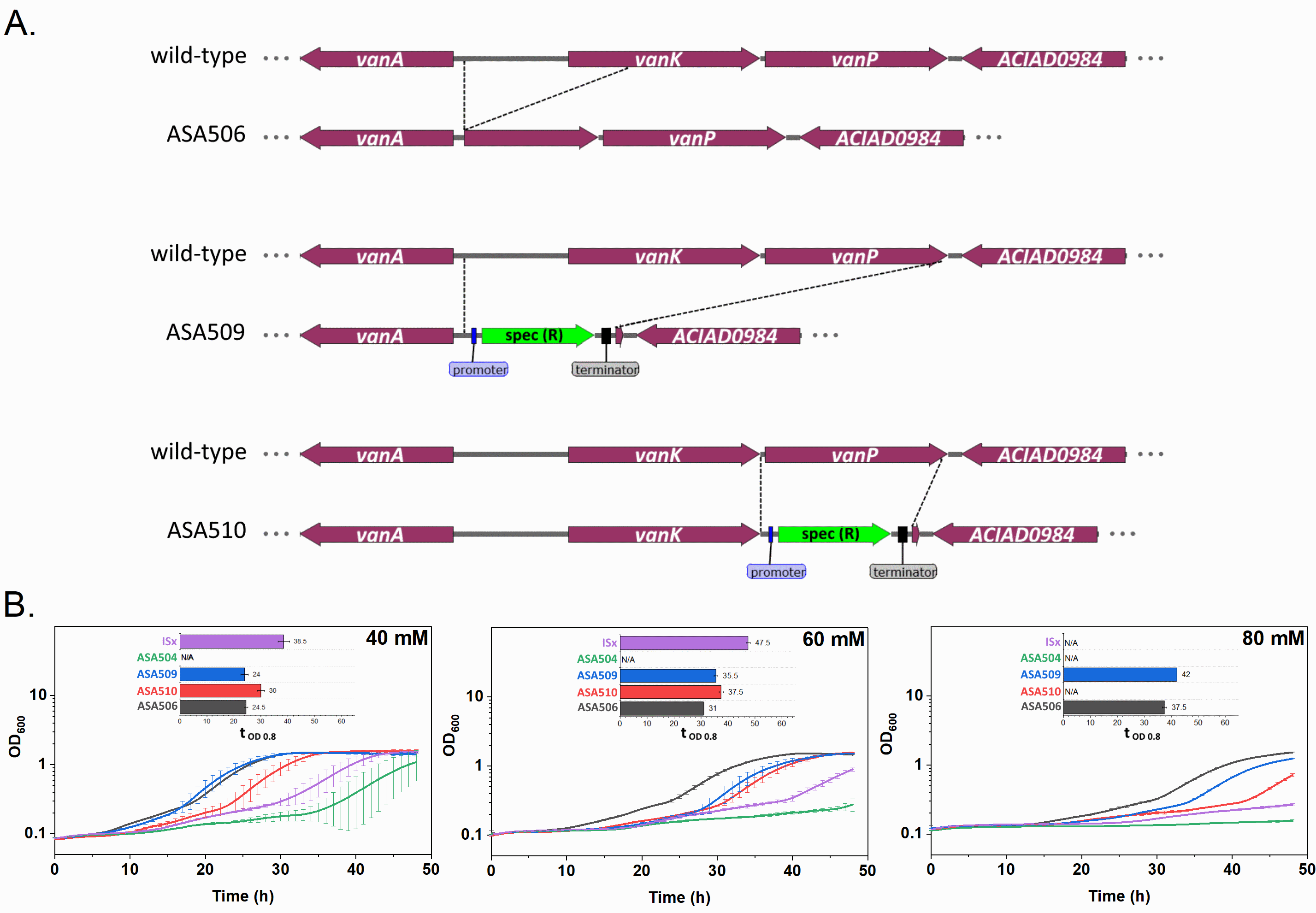
**Figure S10.** Growth comparison between ASA506, ASA509, and ASA510 on vanillate. (A) Genomic context of the *vanK*/*vanKP*/*vanP* deletion region in the strains studied. ASA506 was constructed by RAMSES, and ASA509 amd ASA510 were constructed using the knock-out cassettes containing the spectinomycin resistane gene. (B) Growth curves of the strains tested in 40, 60 mM, and 80 mM vanillate. ISx and ASA504 was used as the controls. All the strains were cultivated in mineral salts media with vanillate as the sole carbon source. Time spent for the cells to reach the OD of 0.8 was used as the indicator to evaluate the tolerance (t_OD 0.8_). The indicator was calculated only when both replicates reached OD 0.8 within 48 h. All the values were calculated from two replicates and the error bars indicate the standard deviations. The y-axis is shown in log10 scale.

**References**

1.	Luo J, Lehtinen T, Efimova E, Santala V, Santala S. 2019. Synthetic metabolic pathway for the production of 1-alkenes from lignin-derived molecules. Microb Cell Fact 18:48.

2.	Suárez GA, Renda BA, Dasgupta A, Barrick JE. 2017. Reduced Mutation Rate and Increased Transformability of Transposon-Free *Acinetobacter baylyi* ADP1-ISx. Appl Environ Microbiol 83.

3.	Nichols NN, Harwood CS. 1997. PcaK, a high-affinity permease for the aromatic compounds 4-hydroxybenzoate and protocatechuate from *Pseudomonas putida*. J Bacteriol 179:5056–5061.

4.	Smith MA, Weaver VB, Young DM, Ornston LN. 2003. Genes for Chlorogenate and Hydroxycinnamate Catabolism (*hca*) Are Linked to Functionally Related Genes in the *dca-pca-qui-pob-hca* Chromosomal Cluster of *Acinetobacter* sp. Strain ADP1. Appl Environ Microbiol 69:524–532.
